# Pharmacological and behavioral effects of tryptamines present in psilocybin-containing mushrooms

**DOI:** 10.1101/2023.10.19.563138

**Authors:** Ryan J. Rakoczy, Grace N. Runge, Abhishek K. Sen, Oscar Sandoval, Quynh Nguyen, Brianna R. Roberts, Jon H. Sciortino, William J. Gibbons, Lucas M. Friedberg, J. Andrew Jones, Matthew S. McMurray

## Abstract

Demand for more efficacious antidepressants, particularly those with a rapid onset of action, has resulted in a reevaluation of psychedelic drugs for their therapeutic potential. Several tryptamines found in psilocybin-containing ‘magic’ mushrooms share chemical similarities with psilocybin, and early work suggests they may also share receptor targets. However, few studies have explored their pharmacological and behavioral effects. To accomplish this, we compared baeocystin, norbaeocystin, and aeruginascin with psilocybin to determine if they are metabolized by the same enzymes, penetrate the blood brain barrier, serve as ligands for similar centrally located receptors, and modulate behavior in rodents similarly. We first assessed the stability and optimal storage and handling conditions for each compound. *In vitro* enzyme kinetics assays then found that all compounds shared nearly identical rates of dephosphorylation via alkaline phosphatase and metabolism by monoamine oxidase. Further, we found that only the dephosphorylated products of baeocystin and norbaeocystin could cross a blood brain barrier mimetic to a similar degree as the dephosphorylated form of psilocybin, psilocin. Behaviorally, only psilocybin was found to induce head twitch responses in rats, a marker of 5HT2A agonism and indicator of the compound’s hallucinogenic potential. However, like psilocybin, norbaeocystin was also found to improve outcomes in the forced swim test. All compounds were found to cause minimal changes to metrics of renal and hepatic health, suggesting innocuous safety profiles. Collectively, this work suggests that other naturally-occurring tryptamines, especially norbaeocystin, may share the same therapeutic potential as psilocybin, but without causing hallucinations.

**HIGHLIGHTS:**

- Baeocystin, norbaeocystin, and aeruginascin may have similar therapeutic value to psilocybin, but are understudied
- Compound stability varied widely, with dephosphorylated forms showing lowest stability
- Rates of metabolism by alkaline phosphatase and monoamine oxidase were similar across compounds
- Blood brain barrier penetration was limited to dephosphorylated forms of psilocybin, baeocystin, and norbaeocystin
- Rat behavioral testing suggested norbaeocystin may have therapeutic utility similar to psilocybin, without causing hallucinations

## INTRODUCTION

Demand for more efficacious mental health treatments has resulted in a reevaluation of psychedelic drugs for their therapeutic potential, with recent emphasis on tryptamine alkaloids produced by “magic” mushrooms. Psilocybin, the primary psychoactive tryptamine alkaloid in these mushrooms, has been demonstrated to ameliorate symptoms of several mental health disorders in humans (Johnson et al., 2014; Carhart-Harris et al., 2017; Carhart-Harris et al., 2021). Outcomes such as these have reinvigorated research aimed at identifying, producing, and characterizing other tryptamines, both naturally occurring and synthetic, with potential clinically utility (Blei et al., 2018; Adams et al., 2019; Flower et al., 2023). Such investigations have identified and begun characterizing several psilocybin-related tryptamines also produced in “magic” mushrooms: aeruginascin, baeocystin, and norbaeocystin (Leung & Paul, 1968; Repke & Leslie, 1977; Jacob & Shulgin, 1994; Shulgin & Shulgin, 1997; Sherwood et al., 2020; Glatfelter et al., 2022; Plazas & Faraone, 2023). These studies suggested that these tryptamines may exhibit similar pharmacodynamics to psilocybin and psilocin (the dephosphorylated and more psychoactive form of psilocybin), implying they may also have clinical utility. For example, prior *in vitro* work found that the dephosphorylated form of baeocystin, norpsilocin, has appreciable affinity for several isoforms of serotonin receptors which include many of the same molecular targets as psilocin (Chadeayne et al., 2020; Sherwood et al., 2020; Glatfelter et al., 2022). However, systematic assessments of the complex pharmacologies of these compounds have yet to be completed. Likewise, experiments testing the therapeutic efficacy of these tryptamines in pre-clinical models have not yet been performed. Further research with these compounds may provide insight into a shared mechanism of action for a broad array of psychedelic compounds and could help to identify novel treatments using these and other tryptamines.

Due to their structural similarity, foundational pharmacological assessments of these understudied compounds should be guided by what is known about psilocybin and psilocin. To be capable of exerting psychoactive effects, canonical theory suggests that psilocybin and psilocin act on centrally located targets (e.g., 5-HT receptors) to mediate their behavioral and therapeutic effects. Quickly after ingesting “magic” mushrooms, for example, psilocybin is enzymatically dephosphorylated by alkaline phosphatase, converting the compound from its prodrug form to its more active form, psilocin (Horita & Weber, 1961a; Horita & Weber, 1961b; Horita, 1963; Dinis-Oliveira, 2017). With increased lipophilicity, psilocin is capable of passively crossing the blood brain barrier (BBB) where it interacts with central targets (i.e., 5-HT receptors) and exerts psychoactive effects (Horita & Weber, 1962). Psilocin is then metabolized by monoamine oxidase-A (MAO-A) into comparatively inert metabolites (Passie et al., 2002).

Prior work speculates that other tryptamines present in psilocybin-producing mushrooms may also require dephosphorylation to facilitate membrane permeability (Passie et al., 2002; Dinis-Oliveira, 2017; Zohairi, Khandelia, & Zanjani, 2023), but this has not been confirmed experimentally. Demonstrating that these tryptamines are dephosphorylated via alkaline phosphatase would provide insight into their activation dynamics and additional confirmation of analogous pharmacology. Further, confirming that the dephosphorylated forms of these compounds can passively permeate a biological membrane would provide evidence of their ability to enter the brain wherein they may exert psychoactive effects (Clemons, Kretsch, & Verbeck, 2014). Additionally, it has not yet been determined if the active, dephosphorylated forms of these tryptamines are deaminated by monoamine oxidase A (MAO-A), like psilocin. Since the psychoactive effects of psychedelic tryptamines (i.e., DMT, psilocin) conclude once they are metabolized by MAO-A (Horita, 1963; Blei et al., 2020), characterizing the enzymatic metabolism of other tryptamines is paramount to understanding the pharmacology of these compounds. Furthermore, if these tryptamines are substrates for MAO-A, they may compete with psilocin for binding *in vivo* when consumed in conjunction with psilocybin (i.e., in “magic” mushrooms). This competitive action could potentially increase psilocin’s half-life and augment its psychoactivity (Blei et al., 2020). At minimum, studying the enzymatic dephosphorylation and deamination kinetics associated with each of these tryptamines is necessary before more comprehensive clinical assessments can be conducted.

In addition to biochemical studies, these psilocybin-related tryptamines may also be capable of exerting quantifiable effects on behavior. Specifically, if these compounds are effective in modifying unconditioned behaviors indicative of their purported psychoactive effects, further studies would be warranted to explore their potential therapeutic application. Psilocybin’s effect on rodent head twitch responses has been the most thoroughly studied behavioral effect and is known to be dependent on the activation of 5-HT2A receptors. Head twitch responses (e.g., wet dog shakes) have been established as a reflection of psilocybin’s hallucinogenic nature (Corne & Pickering, 1967; González-Maeso et al., 2007), but considerable debate regarding whether hallucinations are necessary for the therapeutic effects of psilocybin (Hesselgrave et al., 2021; Cao et al., 2022). Additionally, psilocybin’s effects on measures of depression-associated behaviors have been well-studied in rodent models (Hibicke et al., 2019; Hibicke et al., 2020; Hesselgrave et al., 2021). Classically, improvements in outcomes measured by the forced swim test have been associated with strong therapeutic efficacy in humans (Lucki, 1997; Yuen, Swanson, & Witkin, 2017). although other rodent tests of therapeutic efficacy exist but vary by disease-relevance. Sharing any or all of these behavioral effects would support further testing of these compounds as potential antidepressants. Furthermore, to be candidate therapeutics, *in vivo* toxicological assessment of these tryptamines should parallel that of psilocybin, which has the most innocuous safety profile of any classical psychedelic and is well-tolerated by patients in clinical settings (Gable, 1994; Gable, 2004; Hendricks, Johnson, & Griffiths, 2015). Specifically, blood panel testing for markers of renal and hepatic health in rodents administered these tryptamines is necessary to determine the safety and suitability of these compounds for future clinical studies.

Thus, the goal of this work is to provide insights into the pharmacological, behavioral, and toxicological effects of the psilocybin-related tryptamines: aeruginascin, baeocystin, and norbaeocystin. Previous research speculates about the pharmacology of each of these compounds based on their chemical structure similarity to psilocybin, but few studies have investigated the basic assumptions made about these compounds. Further biochemical characterization is essential and may identify novel compounds with clinical utility. To accomplish this, we first synthesized these tryptamines and evaluated their thermal stability to account for sample degradation in the following studies. Next, we determined their relative rates of metabolism by alkaline phosphatase and MAO-A, as these would dictate both their likely site of action (brain vs periphery) and regulate duration of action. Then, we evaluated the potential of prodrug and dephosphorylated forms of each compound to passively permeate a blood brain barrier mimetic and quantified affinities for interaction with a variety of central receptors. Having determined these attributes, we finally assessed the relative safety of each compound and their efficacy in modulating unconditioned behavioral responses through forced swim testing and observations of head twitch responses in rats. Collectively, this body of work establishes a foundation for assessing the pharmacodynamics of these understudied tryptamines with an emphasis on their potential clinical use (Figure 1).

**Figure 1:**
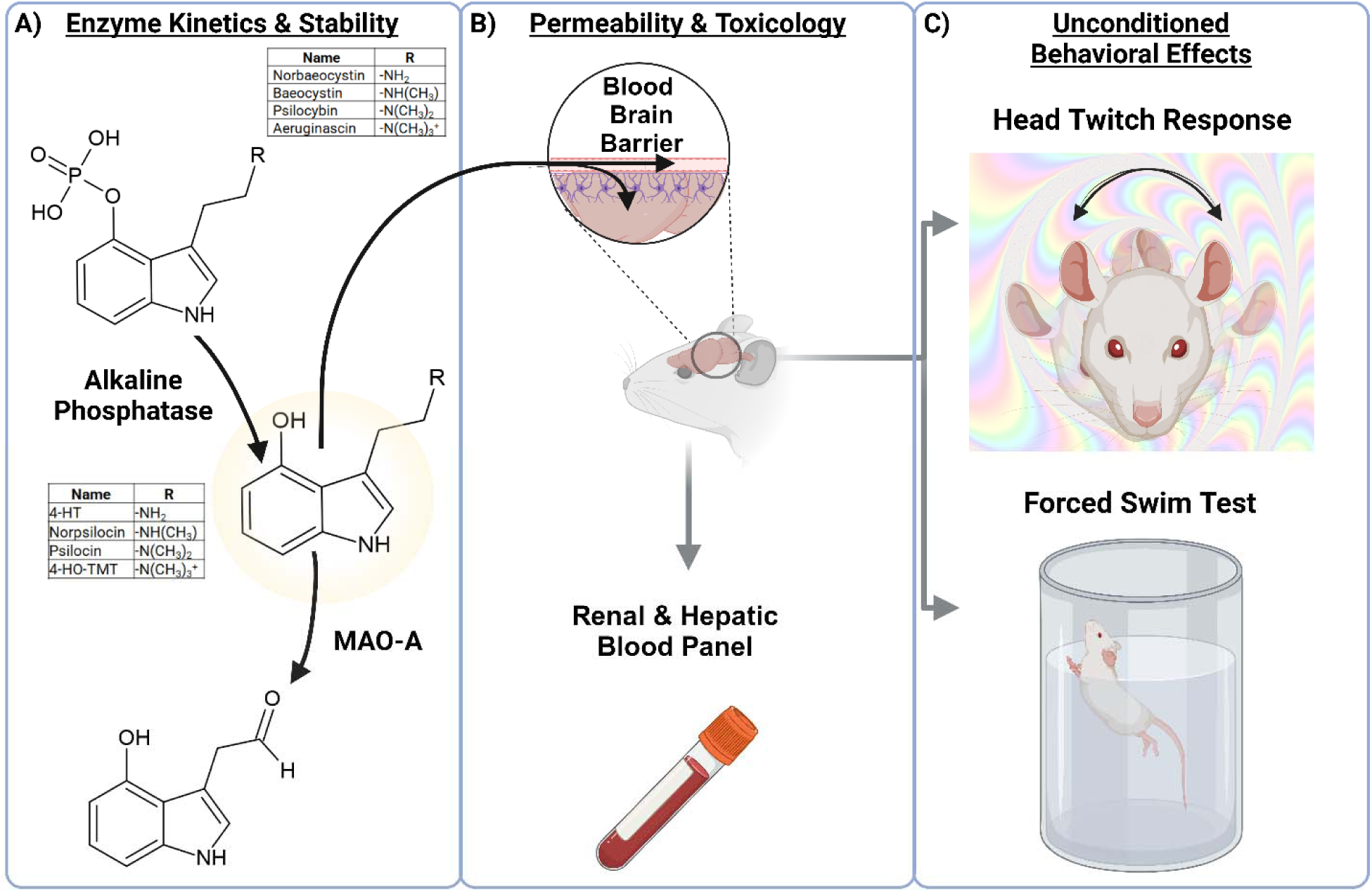
Overview of experiments performed to characterize the pharmacology of psilocybin and the related tryptamines naturally occurring in “magic” mushrooms: norbaeocystin, baeocystin, and aeruginascin. A) Stability of the compounds was tested, followed by determining whether the proforms and dephosphorylated forms were substrates for alkaline phosphatase and monoamine oxidase A, respectively. B) Passive permeability across a blood brain barrier mimetic was determined for proforms and dephosphorylated forms, and renal and hepatic blood panels conducted after proforms were administered to rats. C) The ability of these compounds to modify unconditioned behaviors (head twitch response and forced swim test) was then assessed. Note: this figure was generated using BioRender (www.biorender.com).

## METHODS

### Tryptamine Production, Storage, and Stability Analysis

The tryptamines psilocybin, baeocystin, and norbaeocystin used in this work were biosynthesized using engineered recombinant *E. coli* BL21(DE3) as previously reported (Adams et al., 2019; Adams et al., 2022). Norbaeocystin was purified independently, while baeocystin and psilocybin were purified from the same fermentation broth and separated via preparative HPLC (Adams et al., 2022). The final product was freeze dried, resulting in a white (psilocybin) or light brown (baeocystin and norbaeocystin) powder which was resolubilized in water (∼ 3.52 mM) and passed through an endotoxin removal filter (Pall Corporation - Acrodisc® Unit, MSTG25E3). The resulting endotoxin free solution was concentrated using rotary evaporation followed by freeze drying. Final endotoxin levels were measured using a commercial endotoxin quantification kit (Pierce^TM^ Chromogenic Endotoxin Quant Kit, A39552) and levels were confirmed to be below the limit of detection (< 1 EU/mg).

Aeruginascin synthesis (1.0 g scale, 90% yield) followed published traditional chemical synthesis methods (Sherwood et al., 2020), beginning with purified psilocybin derived from the biosynthetic process described above. The resulting aeruginascin product was purified using preparative HPLC. The collected fractions were concentrated by rotary evaporation, freeze dried, and endotoxin was removed as described above. All products were stored as dry powders at −20 °C.

The thermal stability of the four prodrugs under investigation in this study was evaluated in an aqueous solution at neutral pH, and concentrations ranging between 0.5-5 mM. Four temperatures (4 °C, 37 °C, 60 °C, and 80 °C) were chosen for the analysis of psilocybin, with all four compounds being evaluated at 4 °C and 80 °C. Samples in a broth vehicle were tested in unpurified form in spent cell culture media with cell biomass removed, while the pure components were evaluated in MilliQ. The length of the studies was dependent on temperature, with the 80 °C studies concluding in roughly 1 week, while the 4 °C studies continued for roughly 9 months.

The thermal stability of the four active (dephosphorylated) forms under investigation in this study was evaluated in an aqueous solution at neutral pH, and concentrations ranging between 0.5-5 mM. The active forms were generated through the addition of commercially available Alkaline Phosphatase (AP, Anza^TM^ ThermoFisher IVGN2208; [40 U/mL]) to each prodrug to catalyze the conversion of prodrug to active form. Briefly, a ∼6 mM drug solution in Tris buffer (per 100 mL: 0.58 g NaCl, 0.10 g MgCl, 1.2 g Tris, pH 7.4) is combined with 10 uL/mL AP for 30 min at 37 °C and then the AP is denatured at 80 °C for 20 min before the start of the thermal stability study. Reactions were generally performed at 0.2 mL scale with the temperature controlled using a programmable PCR thermal cycler.

Four temperatures (4 °C, 20 °C, 37 °C, and 60 °C) were chosen for the analysis of psilocin, and all four compounds were evaluated at 37 °C. Samples were evaluated in the Tris AP buffer detailed above. The length of the studies was dependent on temperature, with the 60 °C studies concluding in roughly 5 hours, while the 4 °C studies continued for 72 hours. All samples were quantified by HPLC.

HPLC-MS analysis was performed as previously reported (Adams et al., 2022). Briefly, samples were prepared by centrifugation at 21,000 rcf for 5 minutes; 2 µL of the resulting supernatant was then injected for analysis. Analysis was performed on a Thermo Scientific Ultimate 3000 High-Performance Liquid Chromatography (HPLC) system equipped with Diode Array Detector (DAD) and Thermo Scientific ISQ^TM^ EC single quadrupole mass spectrometer with a heated electrospray ionization source, operated in positive mode. Metabolite separation was performed using an Agilent Zorbax Eclipse XDB-C18 analytical column (3.0 mm x 250 mm, 5 µm) with mobile phases of water (A) and acetonitrile (B), both containing 0.1% formic acid at a rate of 1.0 mL/min: 0 min, 5% B; 0.43 min, 5% B; 5.15 min, 19% B; 6.44 min, 100% B; 7.73 min, 100% B; 7.73 min, 5% B; 9.87 min, 5% B. This method resulted in the following observed retention times as verified by analytical standards (when commercially available): norbaeocystin (1.5 min), 4-hydroxytryptamine (3.3 min), psilocybin (2.3 min), psilocin (4.2 min), baeocystin (1.9 min), norpsilocin (3.59 min), aeruginascin (1.85 min) and 4-hydroxy-*N,N,N*-trimethyltryptamine (4-OH TMT, 4.4 min).

### Alkaline Phosphatase Enzyme Kinetics

Once ingested, phosphorylated tryptamines are rapidly hydrolyzed into their active forms via alkaline phosphatase (AP). Pilot experiments first determined compounds were substrates for AP (Figure S1). AP-mediated dephosphorylation rates were then quantified for psilocybin, aeruginascin, norbaeocystin, and baeocystin by measuring their conversion into their respective active forms: psilocin, 4-hydroxy-*N,N,N*-trimethyltryptamine (4-HO-TMT), 4-hydroxytryptamine (4-HT), and norpsilocin, respectively (Horita & Weber, 1961a Horita & Weber, 1961b; Sherwood et al., 2020; Klein et al., 2021; Glatfelter et al., 2022). One millimolar tryptamine and AP-enzyme (ThermoFisher IVGN2208; [40 U/mL]) stock solutions were prepared in AP-buffer (100 mM NaCl, 5 mM MgCl2, 100 mM Tris-HCl, to 7.4 pH with NaOH), aliquoted in 1.5 mL centrifuge tubes and stored at −20 °C until use. On the day of experiments, tryptamine and AP-enzyme stocks were thawed, and reaction mixtures prepared (100 µL total volume) in plastic, glass-vial inserts for HPLC auto-sampling (ThermoFisher 6EME03CPPSPT) by diluting stock substrates in AP-buffer to the following concentrations (in µM): 10, 25, 33, 50, 100, 150, 200, 300. Prior to addition of enzyme, baseline readings were determined by injecting 2 µL of the reaction mixture into the HPLC as described above (Adams et al., 2022). All data was managed and processed using Thermo Scientific Chromeleon 7.3 Chromatography Data Management System. After baseline readings, 2 µL of AP-enzyme (reaction concentration: [0.8 U/mL]) was added to the reaction tube and 2 µL of the reaction mixture auto-sampled every 10-minutes for one-hour while being held at room temperature (data in Figure S2). Duplicate reactions were run for each substrate concentration. The peak area derived from UV chromatographs for each dephosphorylated tryptamine was divided by the baseline peak area of the unmodified substrate plus the area of the product to obtain conversion rates (Hong, Zhang, & Lin, 2018). Conversion rates were normalized to time to calculate initial velocities for the reaction (Dean, 2000; Hong, Zhang, & Lin, 2018). GraphPad Prism (v8.4.3) was then used to fit the initial velocities for each substrate concentration to the Michaelis-Menten equation to obtain V_max_ and K_m_ for each substrate.

### Monoamine Oxidase Enzyme Kinetics

Monoamine oxidase A (MAO-A) deamination of psilocin and related, dephosphorylated tryptamines, in part, mediates the termination of their psychoactive effects through conversion of the active forms of compounds into inert metabolites, such as 4-hydroxyindole-3-acetaldehyde and 4-hydroxyindole-3-acetic acid (Horita, 1963; Blei et al., 2020). A byproduct of this enzymatic conversion is the generation of hydrogen peroxide, which was used to indirectly quantify rates of MAO-A-mediated conversion of the dephosphorylated, active forms of the tryptamines.

Immediately before beginning the experiments, 1 mM stock of dephosphorylated tryptamines in PBS (Millipore 6505) were generated by incubating tryptamines for 30-minutes at 37 °C with alkaline phosphatase (ThermoFisher IVGN2208; 20 U/mL), which was inactivated immediately after incubation via heating to 80 °C for ten-minutes, and stock solutions then stored on ice until use. 1 mM 5-HT (Serotonin HCl; Sigma H9523) was prepared and incubated with enzyme identically as the rest of the tryptamines. A commercially available kit (Abcam ab138886) was then used to fluorometrically quantify hydrogen peroxide levels and was prepared according to manufacturer’s protocol. Briefly, each well in a 96-well microplate was filled with 50 µL of the kit-provided reaction mixture, and duplicate wells were filled with 40 µL of each dephosphorylated tryptamine diluted to achieve the tested concentrations (i.e., 500, 250, 100, 50, & 33 µM) immediately before use. Kit-provided hydrogen peroxide standards were tested in duplicate on the same microplate and used to generate a standard curve. Next, 10 µL of previously diluted MAO-A (Corning Supersomes FisherSci NC9906032) was added to wells containing dephosphorylated tryptamines to achieve a final concentration of 0.05 mg/mL. Duplicate control wells contained [500 µM] tryptamines and were incubated without MAO-A to control for any spontaneous H_2_O_2_ generation. The microplate was then incubated and allowed to equilibrate at 37°C in a microplate reader (BioTek Cytation 5) while kinetic fluorescence readings (ex./em.: 640/680) were taken of each well every 2-minutes over 55-minutes (data in Figure S3). Hydrogen peroxide concentrations were calculated using the standard curve generated in the same assay plate according to the assay protocol. Graphpad Prism software was used for analysis. Rates of generation of H_2_O_2_ were calculated by normalizing H_2_O_2_ concentrations to time for each concentration and these velocities were fit to the Michaelis-Menten equation using GraphPad Prism to obtain V_max_ and K_m_ for each tryptamine.

### Membrane Permeability

A parallel artificial membrane permeability assay blood brain barrier kit (Corning PAMPA 353015) was used to determine which tryptamines (both the proforms and “active”, dephosphorylated forms) were capable of passively permeating a blood brain barrier (BBB) mimetic (Chen et al., 2008; Clemons, Kretsch, & Verbeck, 2014). Following manufacturer’s protocol, identical concentrations (200 µM in PBS) of psilocybin, psilocin, norbaeocystin, 4-hydroxytryptamine (4-HT), baeocystin, norpsilocin, aeruginascin, 4-hydroxy-N,N,N- trimethyltryptamine (4-HO-TMT), and 5-HT (Serotonin HCl; Sigma H9523) were placed into a “donor” well (300µL) and separate permeability control acceptor wells, in triplicate, on a 96-well microplate containing a semi-permeable, lipid-coated membrane that separated compounds from an “acceptor” well filled with an identical buffer (PBS, 7.4 pH, room temperature). “Active” forms of each compound (i.e., psilocin) were generated immediately prior to the experiment using alkaline phosphatase, as described above. The microplate was then incubated for 4 hours at room temperature after which the buffer in the “acceptor” and permeability control acceptor wells was collected (300 µL). Samples were analyzed via HPLC and microplate reader (Agilent Epoch Biotek) using UV absorbance at 280nm to confirm and quantify the concentration and rate at which compounds permeated, respectively (Kansy, Senner, & Gubernator, 1998). HPLC chromatographs are shown in Figure S4 and S5. Using the peak absorbance for each respective test compound, the Permeability Rate (*Pe*) was quantified using the following calculation: 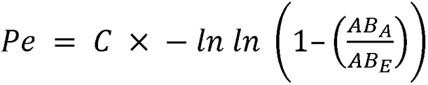. *AB_A_* is the absorbance of solution in the “acceptor” wells minus blank (i.e., PBS, or PBS + heat-inactivated enzyme for “active” forms), *AB_E_* is the absorbance of the “permeability control” wells minus blank and, using a 4-hour incubation, C = 3.47×10^-5^ cm/s. *C* (cm/s) = *V_D_* x *V_A_* (*V_D_* + *V_A_*) x *area* x *time* Where donor well volume (V_D_) is 0.2 cm^3^, acceptor well volume (V_A_) is 0.3 cm^3^, Membrane Area (Area) is 0.24cm^2^, and time is 14,400 sec. Compounds were tested in triplicate and permeability values averaged with standard error of the mean reported. A one-way ANOVA was used to test for significance between the permeability rates of penetrable compounds.

### Identification of Receptor Targets

Receptor binding assays were performed by the NIMH Psychoactive Drug Screening Program (PDSP). Briefly, primary binding assays were used to initially screen compounds at single concentrations (10µM) for the ability to inhibit the binding of a radioligand known to have high affinity for binding with the receptors being tested (i.e., [3H]-Ketanserin for 5HT2A receptors). Compounds with ≥50% radioligand-binding inhibition (i.e., 50% receptor occupancy; IC50 ≤ [10µM]) were next screened in secondary binding assays. Secondary binding assays include testing selected compounds across 11 different concentrations for the ability to inhibit binding of the radioactive ligand to the receptor of interest. Data was used to generate curves for competitive binding and to calculate inhibitory constants (Ki) for each compound. Next, compounds demonstrating a high affinity for receptor targets were tested for their ability to modulate cellular activity through receptor binding. Functional binding assays tested the ability of these compounds to act as agonists or antagonists at the receptors with which they were previously shown to bind. Results demonstrating compounds to achieve ≥30% agonist activity or >50% antagonist activity had secondary experiments performed, generating full dose-response curves to determine potency, efficacy, and IC50 values.

### Blood Analysis Methods

This investigation consisted of 15 male Long Evans rats treated with either psilocybin, norbaeocystin, or baeocystin (n=5/group). Animals received 1 mg/kg of their respective compound via gavage and were left for one hour before the tail nick procedure was performed with blood collected in serum separating tubes. The blood was allowed to clot and then separated in a centrifuge before the serum was transferred to another vial and frozen for storage at −80 °C. 24 hours after drug administration, this procedure was repeated for a total of two blood samples from each animal. All serum samples were analyzed for levels of alkaline phosphatase, total and direct bilirubin, albumin, total protein, creatinine, and blood urea nitrogen (BUN). Results were compared to standard ranges of each of these parameters in rats, provided by VRL Animal Health Diagnostics.

### Rat Head Twitch Response

A total of sixty-seven adult male and female Long Evans rats (Charles River Laboratories, Raleigh, NC) were randomly assigned to one of four treatment groups: psilocybin (n=5 males, 6 females), norbaeocystin (n=9 males, 10 females), baeocystin (n=9 males, 8 females), and aeruginascin (n=10 males, 10 females) and underwent craniotomy surgery (Adams et al., 2022). Briefly, animals anesthetized with isoflurane (2-5% in O_2_) had a neodymium magnet surgically secured to the skull utilizing dental acrylic anchored to screws embedded in the skull that enabled analysis of the head twitch response via a magnetometer (Halberstadt & Geyer, 2013). Characteristic head twitch responses to psychedelics manifest as rapid head and trunk rotational movements that were quantified by analyzing digitized waveform deflections of a magnetic field in a magnetometer recording chamber. Animals were individually housed, given ad libitum access to food and water, remained on a 12:12 hour light cycle, and allowed seven days for post-operative recovery before the start of experiments. Subjects only received administrations of the tryptamine to which they were assigned, at the concentrations: 0, 0.1, 0.2, 1, and 2 mg/kg of psilocybin, norbaeocystin, baeocystin, or aeruginascin. Drugs dissolved in distilled water (vehicle) were administered via intraoral gavage (1.0 mL/kg) and animals were immediately placed in a large polycarbonate tube (∼56 cm diameter, ∼30 cm height). Behavior was recorded for 60 minutes, and the total number of head twitch responses were determined independently by two trained observers. Animals received all concentrations of their designated treatment in a randomized order, with one week separating each drug exposure. All procedures and protocols were conducted in accordance with the National Institutes of Health’s Guidelines for the Care and Use of Laboratory Animals and the Animal Welfare Act and were approved by Miami University’s Institutional Animal Care and Use Committee.

### Rat Forced Swim Test

Methods for the Forced Swim Test (FST) were based on prior a published protocol (Cryan et al., 2005). Briefly, testing occurred over a two-day period, with habitation occurring on day 1 and the FST occurring on day 2. On day 1 (habituation), 60 male Long Evans rats (10 per group) were brought into the procedure room, allowed to rest in their home cages for 30- minutes, and then placed individually into a transparent plastic tube (22 in x 8 in) filled two-thirds of the way (∼16 in) with room temperature (23-26 °C) water. 10 minutes later, animals were removed, dried, and placed in a clean, warmed home cage, and returned to the colony room.

After the habituation, animals were randomly assigned to one of six treatment conditions: vehicle (distilled water), fluoxetine (20 mg/kg), psilocybin (1 mg/kg), norbaeocystin (1 mg/kg), baeocystin (1 mg/kg), or aeruginascin (1 mg/kg). Each treatment was administered via intragastric infusion (1 ml/kg) three times over the following 24-hours: 23.5 hours, 5 hours, and 1 hour before FST testing, matching the published FST protocol (Cryan et al., 2005). After treatment, rats completed the FST in similar conditions to habituation. They were brought into the procedure room, allowed to rest in their home cages for 30-minutes, and then placed in the transparent tubes for only 5 minutes. Behaviors occurring during the FST were video recorded for manual coding of swimming, climbing, and immobility, as described previously (Cryan et al., 2005).

### Rat Behavioral Data Analysis

Two independent observers coded magnetometer data and Forced Swim Test videos, blind to each animal’s treatment condition. Observer scoring was required to match within 90% and was averaged to obtain a final value for that subject for each measure. Final data were analyzed in GraphPad Prism (v.8.4.3), with alpha levels set at 0.05. Head Twitch data were analyzed using separate two-way ANOVAs for each drug, including factors for dose and sex. As prominent and consistent effects of sex were not found, final analyses combined sexes into a single one-way analysis for each drug to increase statistical power. Forced Swim data were analyzed using one-way ANOVA. In both cases, post hoc comparisons were corrected for False Discovery Rate using the two-stage linear set-up method of Benjamini, Krieger, and Yekuteili (Benjamini et al, 2006) was conducted where appropriate to compare vehicle and treatment groups. All analyses were expressed as mean ± SEM.

## Results

### Tryptamine Chemical Characterization

The tryptamine prodrugs synthesized for this study were characterized to confirm identity and purity using LCMS and ^1^H-NMR with water suppression. The analysis of all compounds agreed with expected m/z values (Figures S8C-S11C) and ^1^H-NMR splitting patterns (Figure S12-S15). Purity was determined by ^1^H-NMR (assigned peak area/total peak area) to be: 97.6% - norbaeocystin, 94.6% - baeocystin, 98.8% psilocybin, and 98.6% aeruginascin, peak integration and splitting patterns are provided below. All four prodrugs demonstrated nearly identical UV-Vis spectra with local maximum absorbance at 269 nm (Figure S7B-S11B). HPLC- A280 spectra showed a single symmetric peak with no other significant peaks present (Figures S8A-S11A).

### Thermal Stability

The thermal stability of the produgs (aeruginascin, psilocybin, baeocystin, and norbaeocystin) and their respective activated forms (4OH-TMT, psilocin, norpsilocin, and 4OH- Trm) was evaluated to determine their spontaneous thermal degradation rates under aqueous storage conditions at high (mM) concentrations (Figure S7). In this concentration range, we observed zero order degradation kinetics for all eight compounds with the phosphorylated, prodrug forms demonstrating rate constants 200-600-fold lower than that of the dephosphorylated, active forms. Across the temperatures evaluated, the degree of methylation showed minimal impact on thermal stability. The psilocybin and psilocin degradation rate data demonstrated good fit to the Arrhenius equation across the range of temperatures assessed (Figure S7). Under the conditions tested, the prodrug forms (*e.g.,* psilocybin) were stable at room temperature with minimal degradation (< 10% losses) after weeks of storage, while the active forms (*e.g.,* psilocin) demonstrated appreciable degradation with losses approaching 60% within 24 hours.

### Alkaline Phosphatase Enzyme Kinetics

Rates of AP-mediated dephosphorylation and conversion of the compounds into their “active” forms (Figure 2A) were fit to the Michaelis-Menten equation (curves in Figure 2A; data in Figure S2) and V_max_ and K_m_ calculated for: psilocybin, 0.133 µM min ^-1^ & 109 µM; norbaeocystin, 0.112 µM min ^-1^ & 96.3 µM; baeocystin, 0.135 µM min ^-1^ & 126 µM; and aeruginascin, 0.140 µM min ^-1^ & 133 µM, respectively. A one-way ANOVA determined no significant differences between tryptamines for the rates of AP-mediated dephosphorylation [F(3,28)=0.29, p=0.83]. These rates of alkaline phosphatase-mediated dephosphorylation suggest agnostic enzyme preference for these tryptamines.

**Figure 2:**
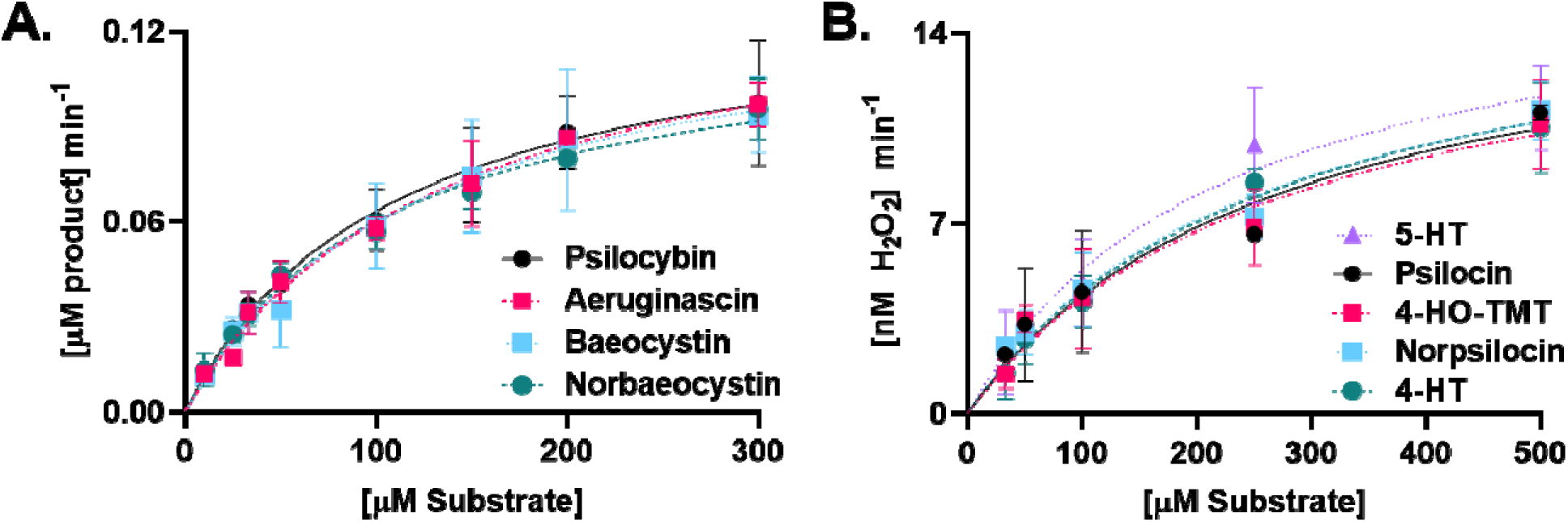
Enzyme kinetics associated with (A) alkaline phosphatase and (B) MAO-A mediated metabolism of tryptamines. No significant differences were noted between compounds in their dephosphorylation rates nor H_2_O_2_ production rates. Results suggest tryptamines of interest are substrates for alkaline phosphatase and MAO-A to a similar degree as psilocybin and psilocin, respectively.

### Monoamine Oxidase Enzyme Kinetics

MAO-A mediated deamination of serotonin (control) and each of the dephosphorylated tryptamines under study was quantified by measuring hydrogen peroxide generation (Figure 2B; data in Figure S3). Production rates of H_2_O_2_ were calculated using the standard curve (R^2^=0.98) and these rates for each tryptamine concentration were fit to the Michaelis-Menten equation (curves on Figure 2B). V_max_ and K_m_ were calculated for: psilocin, 16.3 nM min ^-1^ & 273 µM, 4- HO-TMT, 15.9 nM min ^-1^ & 274 µM, norpsilocin, 16.2 nM min ^-1^ & 253 µM; 4-HT, 16.6 nM min ^-1^ & 269 µM, and 5-HT, 16.7 nM min ^-1^ & 214 µM, respectively. A one-way ANOVA was used to determine there were no significant differences between the rates of hydrogen peroxide production between the tryptamines [F(4, 20)=0.08, p=0.98]. Results are presented as the mean ± the standard error of the mean from duplicate reactions (N=2 for each concentration). Control experiments determined no spontaneous production of H_2_O_2_ in control wells without enzyme.

### Blood Brain Barrier (BBB) Permeability

Both proforms and dephosphorylated, “active” forms of the compounds were tested for their ability to passively permeate an artificial bilipid layer membrane mimicking the BBB (Figure 3, Figure S4, and Figure S5). Compounds capable of permeating the membrane had permeability rates calculated in triplicate and were found to be 1.17×10^-6^ ± 4.21×10^-7^ cm/s for Norpsilocin, 7.77×10^-7^ ± 3.29×10^-7^ cm/s for Psilocin, and 5.58×10^-7^ ± 1.86×10^-7^ cm/s for 4-HT. Neither 4-HO-TMT, 5-HT, nor the proforms of the compounds of interest (i.e., psilocybin) were detected in “acceptor” wells, suggesting no ability to passively permeate the BBB. All compounds were detected in “permeability control” wells and confirmed stable over the length of the assay. A one-way ANOVA found no statistically significant differences between the permeability rates of the detected compounds [F(2, 6)=0.91, p=0.45].

**Figure 3:**
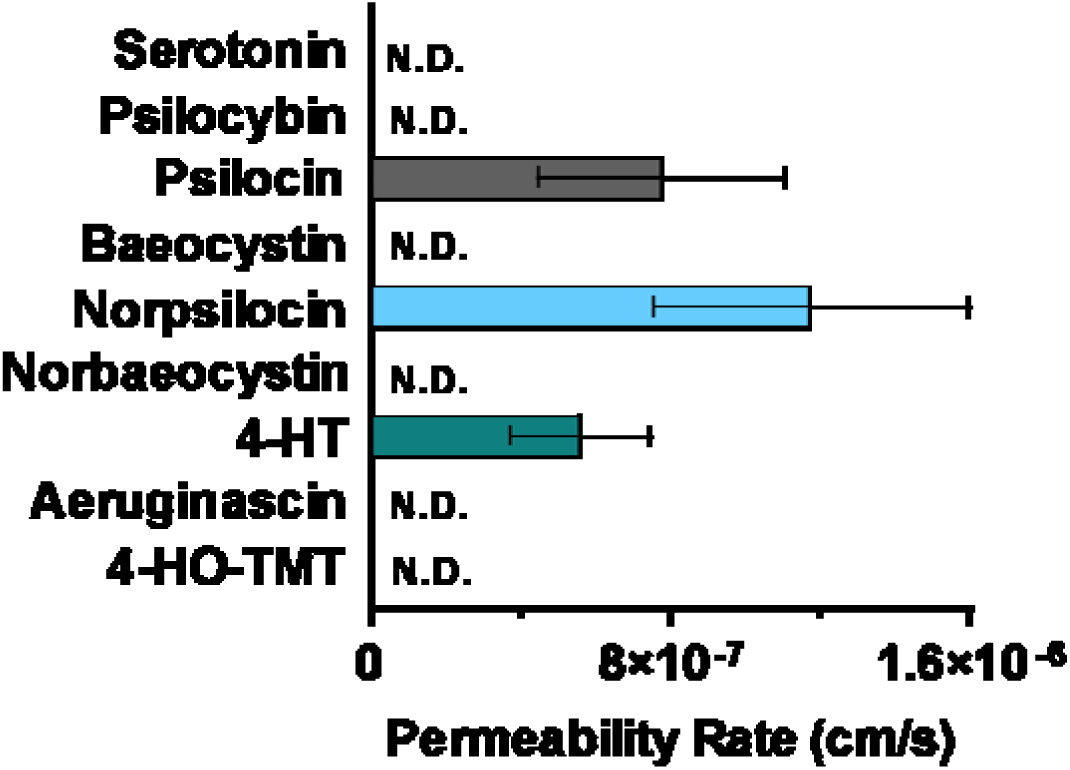
Proforms and dephosphorylated forms of tryptamines [200µM] were assayed in triplicate for their ability to passively permeate a bilipid layer membrane mimicking the blood brain barrier. Psilocin, norpsilocin, and 4-HT were demonstrated to passively permeate the membrane with no significant differences found between their rates (* p≤0.05). Other compounds were not detected (N.D.) in “acceptor” wells, but were confirmed present in control wells via HPLC.

### Blood Analysis

Blood panel results quantifying markers of renal or hepatic health either 1- or 24-hours after receiving psilocybin, norbaeocystin, or baeocystin are shown in Figure 4. Levels of alkaline phosphatase were not significantly greater or less than the normal range for all treatment groups and time points, indicating no abnormalities. Likewise, total and direct bilirubin levels were not significantly greater or less than the normal range for all treatment groups and time points. Albumin levels post-administration of psilocybin at both time points were significantly elevated (1 hour: p=0.04; 24 hours: p=0.04), but total protein levels were not, suggesting a potential change in the albumin/globulin ratio, but not indicative of hepatic or renal damage. A transient rise in serum creatinine and BUN were observed 1 hour after psilocybin administration (p=0.03, and p=0.003, respectively), but returned to within the normal range by 24 hours. Levels of creatinine and BUN stayed within the expected ranges post-administration of norbaeocystin and baeocystin.

**Figure 4:**
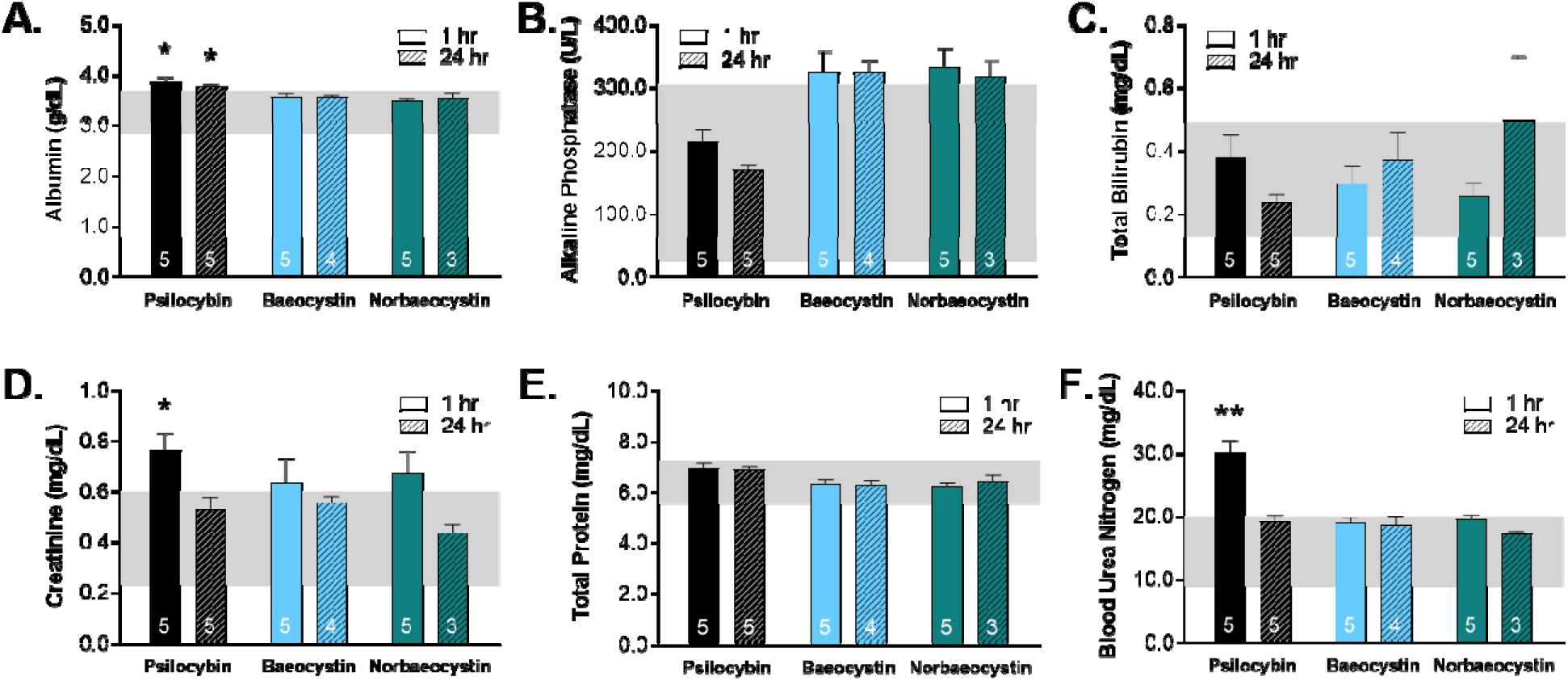
Blood drawn from adult male rats (N=5 per group) 1- or 24-hours post-administration of psilocybin, baeocystin, and norbaeocystin demonstrated animals receiving psilocybin had significantly elevated (A) albumin (at 1- and 24-hours), (D) creatinine (at 1-hour), and (F) blood urea nitrogen levels (at 1-hour); however, (B) alkaline phosphatase, (C) total bilirubin, and (E) total protein levels were all within expected ranges post-administration of any compound. Note: * p≤0.05, ** p≤0.01

### Head Twitch Response Results

Compounds were assessed for their hallucinogenic potential via the head twitch response after animals were administered tryptamines (Figure 5). Animals received treatments of 0, 0.1, 0.2, 1.0, and 2.0 mg/kg of either psilocybin, norbaeocystin, baeocystin, and aeruginascin in water vehicle and head twitch responses recorded for 60-minutes. A one-way ANOVA found a significant effect of psilocybin on head twitch response in both males and females ((F(4,19)=3.54, p=0.03) and (F(4,23)=3.76, p=0.02), respectively). The other tryptamines did not exhibit any significant effects of treatment on head twitch response in either sex (one way ANOVAs p>0.05 for all groups). Results from animals separated by sex are presented in Figure S6.

**Figure 5:**
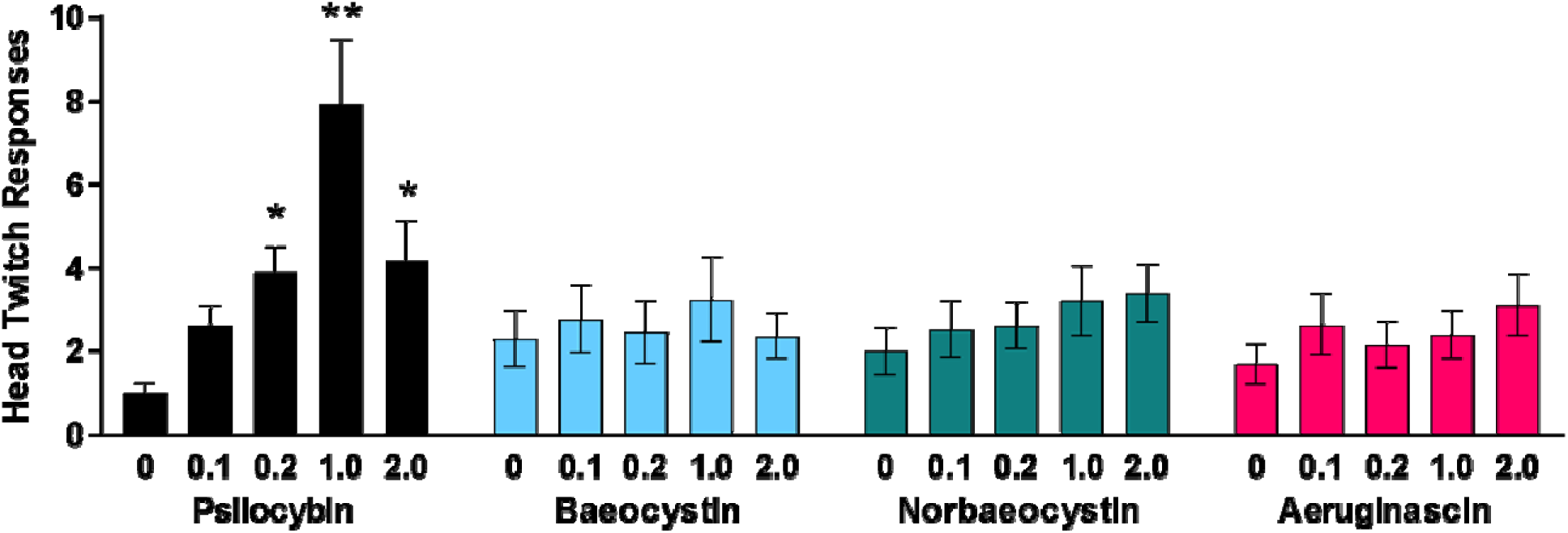
Head twitch responses from male and female rodents receiving varying doses of tryptamines (N=5-10 per sex per group). Compared to vehicle, psilocybin significantly increased HTRs at 0.2, 1.0, and 2.0 mg/kg (* p≤0.05, ** p≤0.01). Baeocystin, norbaeocystin, and aeruginascin were not found to increase head twitches, suggesting they may not have hallucinogenic properties.

### Forced Swim Test

Three behaviors were measured during the FST: immobility, swimming, and climbing (Figure 6). Immobility is thought to be reflective of “defeat”, and is characterized as floating with the absence of significant movement, except that required to keep the rat’s snout above the water. One-way ANOVA found a statistically significant effect of group on immobility [F(5,52)=0.48, p=0.005], with post hoc tests finding psilocybin (p=0.04), norbaeocystin (p=0.05), and fluoxetine (p=0.004) all reduced immobility compared to vehicle (Figure 6A). Swimming assessed active struggling and engagement in the task, and was defined as significant movement of the forepaws, diving to the bottom of the tube, and/or circular movement of the rat in the tube. One-way ANOVA found a significant effect of group [F(5,52)=0.54, p=0.006]. Post hoc tests showed that psilocybin (p=0.04) and fluoxetine (p=0.004) were significantly different from vehicle (Figure 6B). Climbing is reflective of attempts to escape the apparatus, and was defined as movement of the forepaws above the water surface, often paired with strong kicks, which cause the animal to move upwards in the tube. One-way ANOVA found no effect of group [F(5,52)=0.18, p=0.96, Figure 6C].

**Figure 6:**
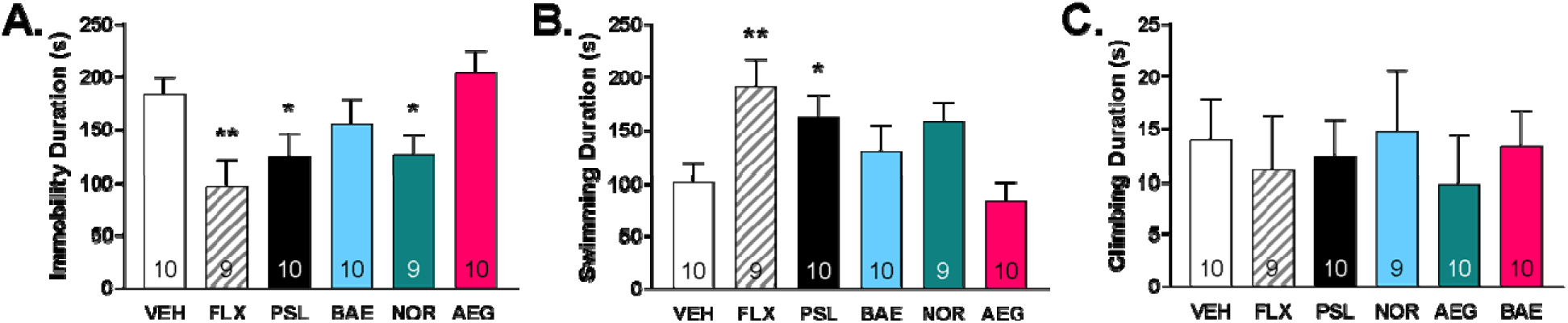
Forced swim test results. Adult male rats (N=10 per group) received 1 mg/kg of fluoxetine (FLX), psilocybin (PSL), norbaeocystin (NOR), baeocystin (BAO), or aeruginascin (AEG). Significant effects on immobility (A) were observed in FLX, PSL, and NOR groups compared with vehicle-treated animals. Increased swimming (B) compared to vehicle receiving animals was noted among FLX and PSL receiving animals. No effects on climbing (C) were observed after any treatment. Note: * p≤0.05, ** p≤0.01

## Discussion

While a significant body of literature has already shed light on the mechanisms and effects of psilocybin, results from the experiments shown here provide some of the first insights into the pharmacological characteristics of aeruginascin, baeocystin, and norbaeocystin. Due to their related chemical structure, these compounds have been speculated to produce similar physiological effects as psilocybin, but thorough biochemical and pharmacological assessments of these tryptamines have not yet been performed. Herein, our thermal degradation assays suggest high stability of all prodrug forms evaluated and results imply extended storage in solution at 4 °C will cause negligible losses in drug potency. In contrast, all active forms showed high degradation rates, even at reduced temperatures, indicating a need to produce these compounds from their prodrug forms on an as-needed basis immediately before use.

Here we show these compounds and their dephosphorylated forms to be substrates for alkaline phosphatase and MAO-A, with similar kinetics to psilocybin and psilocin, respectively. Furthermore, lipophilicity of baeocystin and norbaeocystin was found to be comparable to psilocybin in that the ability of these compounds to cross a bilipid layer is reliant on their dephosphorylation. The unobserved passive permeation of aeruginascin nor its dephosphorylated form, 4-HO-TMT, across a bilipid membrane also supports previous speculation of their inability to cross the BBB (Chue et al., 2022). Receptor binding profiles for aeruginascin, baeocystin, and norbaeocystin, reported here (Supplemental Table 1) and by others, show appreciable affinities for many of the same centrally located targets as psilocybin and psilocin (e.g., serotonergic; Chadeayne et al., 2020; Sherwood et al., 2020; Glatfelter et al., 2022).

Further *in vivo* pharmacological assessment of these compounds tested their ability to modulate unconditioned behaviors that serve as proxies for evaluating their hallucinogenic and antidepressant potential. Specifically, psilocybin was demonstrated to significantly increase HTRs in both male and female rodents, a classic indicator of hallucinogenic activity via central activation of the 5-HT2A receptor (Corne, Pickering, & Warner, 1963; Corne & Pickering, 1967). As norpsilocin and 4-HT (active forms of baeocystin and norbaeocystin, respectively) are evidenced to be capable of crossing the BBB and bind with 5-HT2A receptors, it was surprising that neither induced significant HTRs at any concentration tested. However, these results concur with previous studies demonstrating these compounds to not significantly induce HTRs in rodents (Sherwood et al., 2020; Glatfelter et al., 2022). Forced swim testing demonstrated a comparable effectiveness of psilocybin and norbaeocystin in improving measures related to antidepressant efficacy in rodents. Further, toxicological screening of these compounds determined them to be safe, with profiles comparable to psilocybin when assessing blood markers of renal and hepatic health. Taken together, these investigations into the pharmacology of aeruginascin, baeocystin, and norbaeocystin have highlighted important similarities and differences from psilocybin. Furthermore, these studies have suggested that norbaeocystin may be as effective as psilocybin in improving measures related to antidepressant efficacy. However, unlike psilocybin, norbaeocystin was not found to cause HTRs in rodents and therefore may represent a non-hallucinogenic, therapeutic alternative to psilocybin.

Psychedelic tryptamines exert their hallucinogenic actions via agonism of the 5-HT2A receptor, a G-protein coupled serotonin receptor. Site-specific interactions of psychedelics within the binding pocket of this receptor have been shown to lead to preferential activation of one of several intracellular second messenger signaling cascades with distinct behavioral effects following activation of each pathway. Biased signaling through the β-arrestin or the canonical Gαq-pathway has been demonstrated in experiments using many different psychedelics with high affinity for the 5-HT2A receptor (Kim et al., 2020; Cao et al., 2022; Glatfelter et al., 2022; Poulie et al., 2022). However, it remains unclear if activation of one, or both, signaling pathways mediates the hallucinogenic or antidepressant effects of psychedelics. Previously, it has been suggested that 5-HT2A β-arrestin activity alone is inadequate in producing hallucinogenic effects when assessed via the HTR assay, but may be sufficient in ameliorating symptoms of depression (Cao et al., 2022; Lewis et al., 2023). Furthermore, β- arrestin-2 signaling has been demonstrated to be essential in mediating the therapeutic effects of serotonergic antidepressants (Li et al., 2021). Biased β-arrestin signaling may, in part, explain why tryptamines herein have been shown to have appreciable affinity for the 5-HT2A receptor, but not generate HTRs in rodents. For example, psilocin has been shown to signal through both the Gαq- and β-arrestin pathways (Kim et al., 2020; Glatfelter et al., 2022) and simultaneous activation of these two signaling cascades may lead to production of both HTRs (i.e., hallucinations) and antidepressant effects. Furthermore, biased β-arrestin signaling of the dephosphorylated form of norbaeocystin, 4-HT, may explain its lack of effect on HTRs and efficacy in the forced swim test. An alternative explanation may also lie in recent studies showing that psilocin can directly activate TrkB receptors (Moliner et al. 2023), causing antidepressant effects that are separable from 5HT2A receptor activation. TrkB may be similarly activated by norbaeocystin. Furthermore, the effects of several antidepressant compounds are also not mediated by 5-HT2A activation; the antidepressant actions of ketamine, for example, are mediated through NMDA-receptor antagonism (Krystal, Kavalali, & Monteggia, 2023). Not only does this question the role of 5-HT2A receptors in the antidepressant action of psychedelics, but it highlights a need for further investigating if modulation of non-canonical receptors and pathways by psychedelics mediates their therapeutic effects.

The implications of the results from these experiments should be considered in light of their limitations. Firstly, concentrations of compounds tested for *in vitro* studies were higher than those likely to be reached physiologically after clinical treatment. For example, the therapeutic effects of psilocin may be achieved after reaching only nanomolar concentrations in the brain and blood (Madsen et al., 2019). Thermal degradation kinetics were measured in the [mM] range and it may be the trends observed here do not extend to lower concentrations. Although we observed zero order thermal degradation kinetics in the mM concentration range, preliminary data suggests that these kinetics may transition to first order when concentrations fall into a more biologically relevant range (data not shown). Due to the limits of detection of our analytical equipment, however, we were unable to test such low tryptamine concentrations in our thermal degradation, alkaline phosphatase, and monoamine oxidase assays. Furthermore, enzyme kinetics assays utilized biologically relevant, but non-human sources of enzymes and future studies should confirm results herein utilizing more physiologically relevant enzyme isoforms and conditions. The MAO-A assay also indirectly measured kinetics via H_2_O_2_ evolution, a product of tryptamine deamination. Although this method has been used effectively in similar kinetics experiments and serves as a well-documented proxy for assessing MAO-A deamination rates (Green & Haughton, 1961; Reis & Binda, 2022), it is not a direct measure and does not preclude H_2_O_2_ interaction with other metabolic products. Studies in humans, nonetheless, have previously shown *in vivo* elimination of psilocin to follow 1^st^ order pharmacokinetics and support kinetics results observed herein (Hasler et al., 1997; Holze et al., 2022; Holze et al., 2023). Of importance to mention too is that membrane permeability experiments only assessed the ability of these compounds to passively permeate a blood brain barrier mimetic. For example, results showed that 5-HT did not passively permeate a bilipid layer membrane, but 5-HT is known to be actively transported across the blood brain barrier *in vivo* (Nakatani et al., 2008). Future studies should confirm the results of these enzyme kinetics and blood brain barrier permeability experiments *in vivo*. Lastly, our toxicology work measuring blood markers of renal and hepatic health was only completed in male animals and should be repeated and include females. This was also the case for our tests of antidepressant efficacy, which should be considered preliminary given their reliance on a single behavioral paradigm and exclusion of females. Future studies should confirm results in multiple validated animal models of depression and in females.

In conclusion, these experiments provide one of the first comprehensive pharmacological assessments of the psilocybin-related tryptamines, aeruginascin, baeocystin, and norbaeocystin. When compared alongside psilocybin, these tryptamines (1) exhibited similar kinetic properties as substrates for the enzymes involved in drug activation and degradation pathways, (2) passively permeate a blood brain barrier mimetic (psilocin, 4-HT, and norpsilocin), and (3) serve as ligands for analogous receptors. Behavioral assessments of these compounds’ antidepressant efficacy further identified norbaeocystin as a potential novel and non-hallucinogenic therapeutic. Future studies investigating the pharmacology of the active, dephosphorylated forms of these compounds should take into consideration their rates of degradation when planning experiments with time scales as short as several hours, even at moderately elevated temperatures. Compounds may also be found to exert synergistic pharmacological effects when consumed in combination, for example via “magic” mushrooms. Such “entourage effects” would be of interest to quantify and potentially identify these compounds to be competitive substrates for MAO-A mediating an increased half-life for psilocin *in vivo*. At a minimum, these experiments provide evidence in support of previous conjecture labeling these tryptamines as being “psilocybin-related”, highlight a need for studying non-canonical pathways modulated by psychedelics, and justify further testing of psilocybin and norbaeocystin as potential therapeutics.

## Supporting information

Supplemental Methods and Results

## Acknowledgements

We would like to acknowledge Zoe Platow, Katie Huss, and Kaitlyn Mancini for their assistance with rat behavioral coding (FST and HTR), and C.N. Wyatt and B. Gant for insightful discussions about psychedelics.

