## Supplemental Methods and Results for "Pharmacological and behavioral effects of tryptamines present in psilocybin-containing mushrooms"

**Supplemental Materials for:** Pharmacological and behavioral effects of tryptamines present in psilocybin-containing mushrooms

**AUTHORS:** Ryan J. Rakoczy ^1^, Grace N. Runge ^1^, Abhishek K. Sen ^2^, Oscar Sandoval ^1^, Quynh Nguyen ^2^, Brianna R. Roberts ^1^, Jon H. Sciortino ^1^, William J. Gibbons Jr. ^2^, Lucas M. Friedberg ^2^, J. Andrew Jones ^2^; Matthew S. McMurray ^1^

**Author Affiliations:**

^1^ Department of Psychology, Miami University, Oxford, OH 45056

^2^ Department of Chemical, Paper, and Biomedical Engineering, Miami University, Oxford, OH 45056

**Corresponding Author:**

Matthew McMurray

Assistant Professor

Department of Psychology

Miami University

90 N Patterson Ave

Oxford, OH 45056

513-529-2415

**Supplemental Methods**

Enzyme Mediated Metabolism of Tryptamines

Pilot experiments confirmed the enzyme-mediated metabolism of each tryptamine. Alkaline phosphatase dephosphorylated the pro-forms of the compounds into their active forms which was tracked over time via HPLC and visualized in chromatographs. 500 µM solutions of aeruginascin, baeocystin, norbaeocystin, and psilocybin were prepared in PBS and separated into triplicate vials containing 100 µL of each tryptamine. An initial baseline measurement was performed via HPLC to confirm tryptamine concentration before alkaline phosphatase was added to the vial (8 µL; ~50 EU/mL) and incubated at room temperature for ten minutes. Another measurement was then performed on the HPLC to determine compounds were substrates for alkaline phosphatase. Areas under the curve for each isolated tryptamine peak on UV chromatographs (280 nm) were obtained and compared to control (data not shown). Representative traces from HPLC-derived chromatographs are shown in Figure S1.

**Supplemental Results**

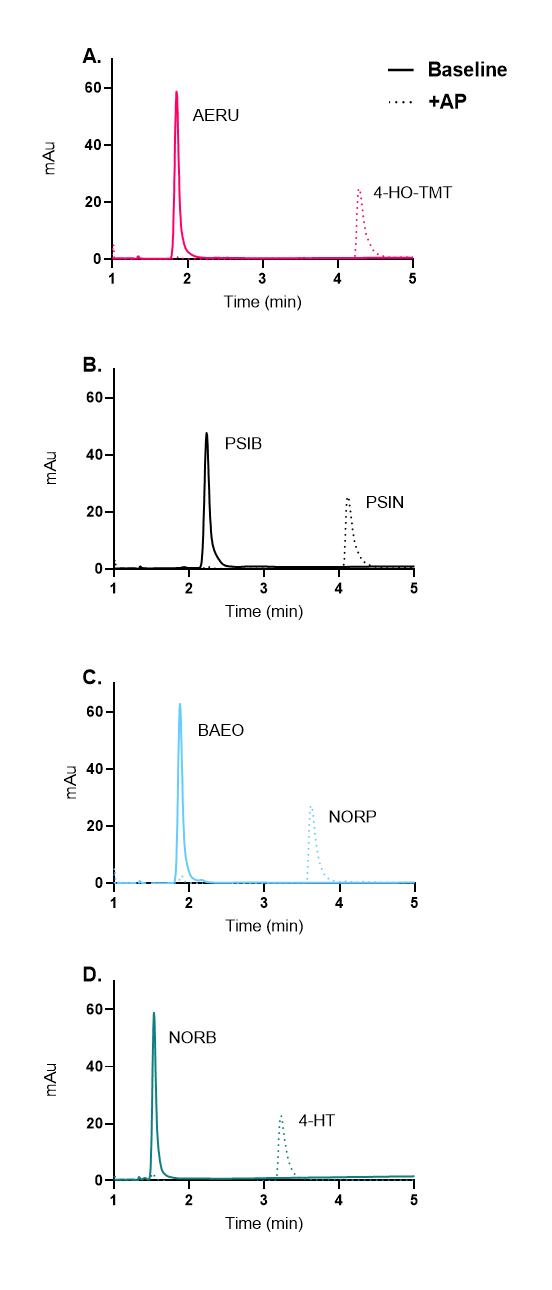

Supplemental Figure S1: Pilot enzyme metabolism experiments suggest alkaline phosphatase to dephosphorylate and convert (A) Aeruginascin (AERU) to 4-HO-TMT, (B) Psilocybin (PSIB) to Psilocin (PSIN), (C) Baeocystin (BAEO) to Norpsilocin (NORP), and (D) Norbaeocystin (NORB) to 4-HT. Chromatograph traces show baseline (before enzyme) and product creation after enzyme addition (+AP) and incubation for ten-minutes at room temperature.

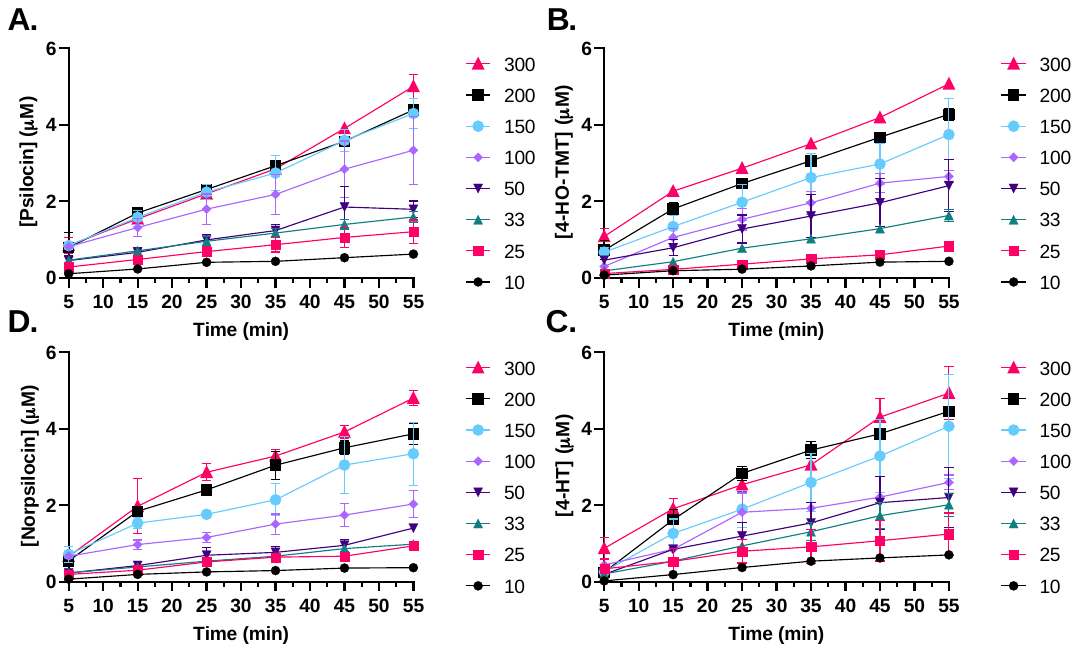

Supplemental Figure S2: Enzyme kinetics data, presented as product formation, for alkaline phosphatase-mediated dephosphorylation of psilocybin (A), aeruginascin (B), baeocystin (C), and norbaeocystin (D) over time. Products were quantified by HPLC. Legend indicates initial concentration (µM) of prodrug in each respective reaction. Data points are the mean ± S.D. from two replicate experiments.

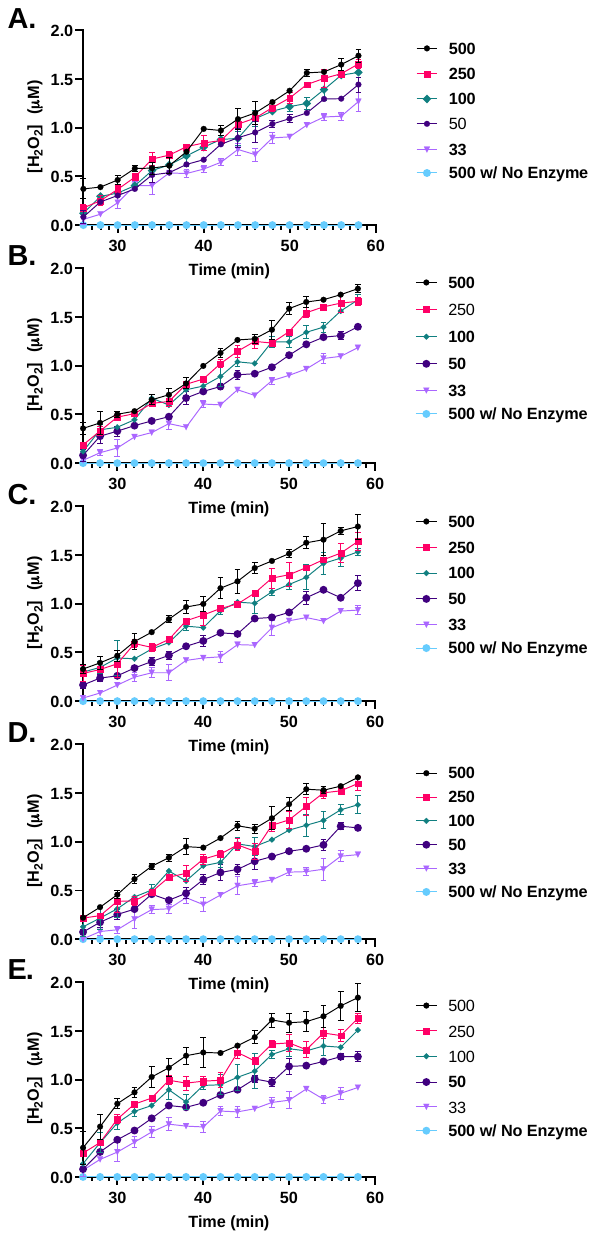

Supplemental Figure S3: MAO-A mediated production of H_2_O_2_ was used as an indirect measurement to quantify metabolism of psilocin (A), 4-HO-TMT (B), norpsilocin (C), 4-HT (D), and 5-HT (E). Control experiments (sans MAO-A) demonstrated no detectable levels of spontaneous evolution of H_2_O­_2_ when [500µM] tryptamines were incubated in the same microplate under identical conditions, but without the addition of MAO-A enzyme (500 µM w/ No Enzyme-A; blue circle). The microplate was equilibrated for 25-minutes at 37°C to with kinetic fluorescence readings taken every 2-minutes for 55-minutes total. Concentrations of substrates in µM and data points are the mean ± S.D. of H_2_O_2_ produced from duplicate wells.

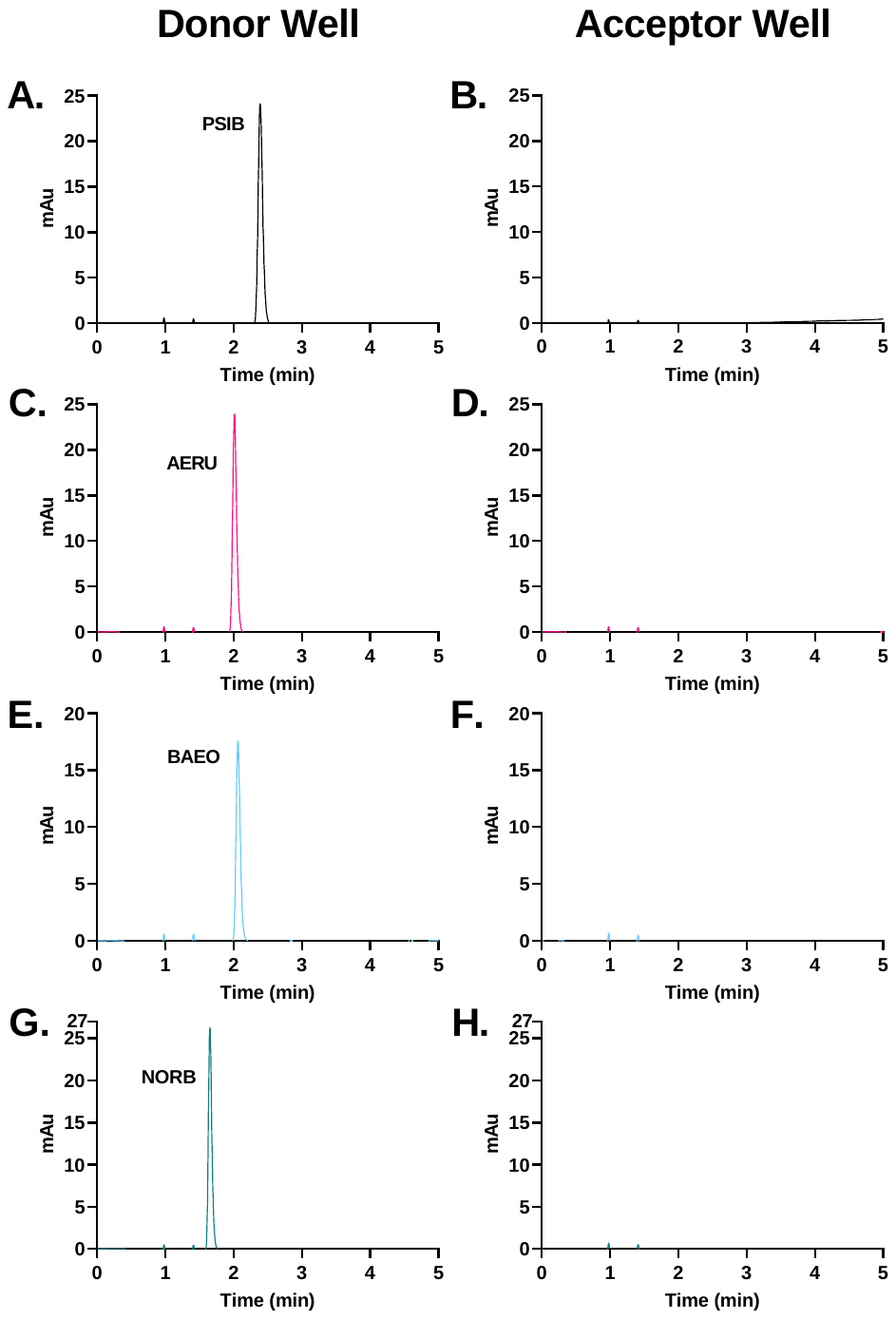

Supplemental Figure S4: Membrane permeability verification HPLC chromatographs. Representative trace of control chromatographs from corresponding donor wells demonstrated (A) psilocybin, (C) aeruginascin, (E) baeocystin, and (G) norbaeocystin were assayed and present at the end of assay incubation via HPLC (left column). Chromatographs (right column) of samples from acceptor wells show (B) psilocybin, (D) aeruginascin, (F) baeocystin, and (H) norbaeocystin were undetected via HPLC and thus not demonstrated to passively permeate the bilipid layer membrane.

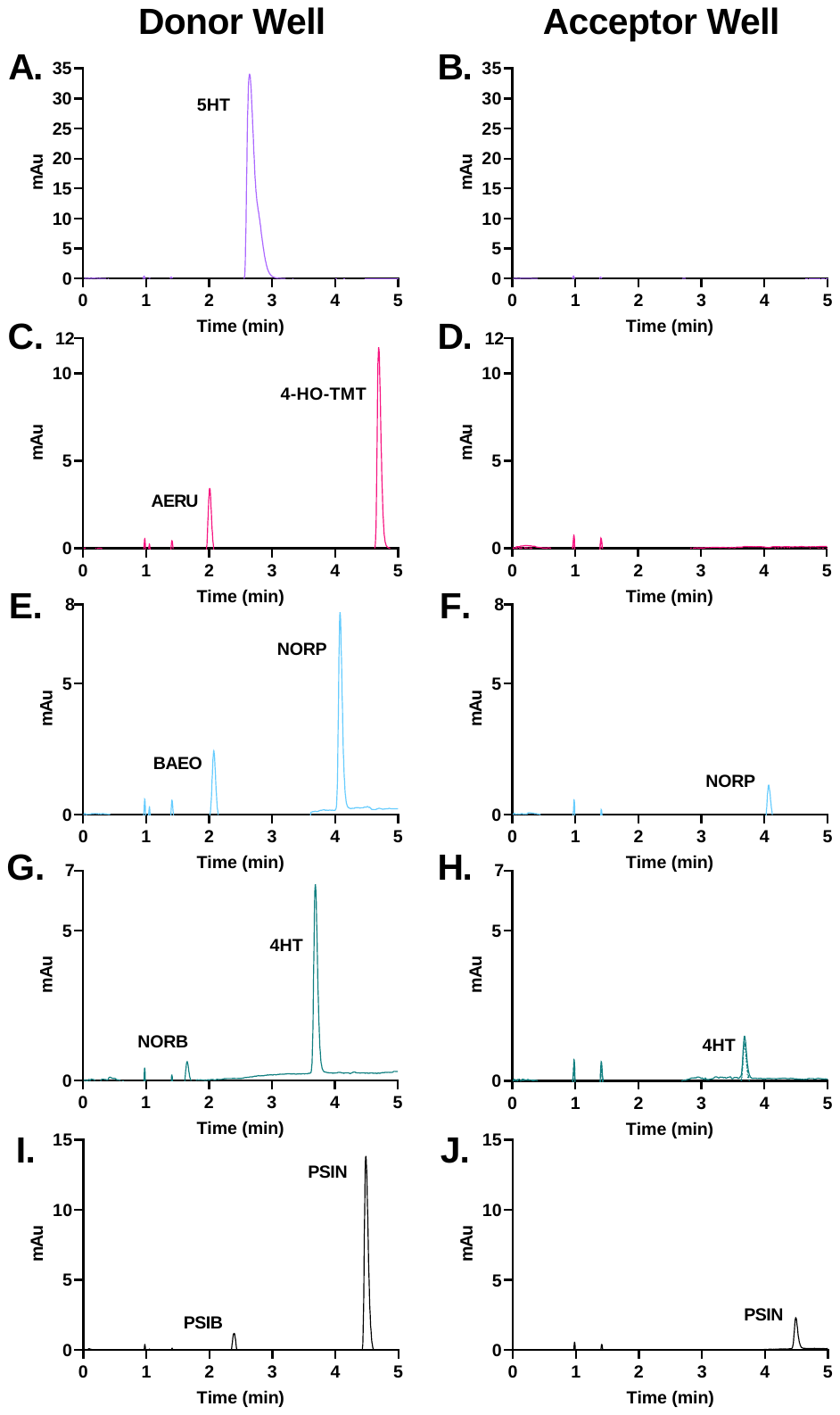

Supplemental Figure S5: Membrane permeability verification HPLC chromatographs. Representative trace of control chromatographs from corresponding donor wells demonstrated (A) 5HT, (C) 4-HO-TMT, (E) norpsilocin, (G) 4HT, and (I) psilocin were assayed and present at the end of assay incubation via HPLC (left column). Chromatographs (right column) of samples from acceptor wells show (F) norpsilocin, (H) 4HT, (F) baeocystin, and (J) psilocin were present and thus capable of passively permeating the bilipid later membrane. Acceptor wells from (B) 5HT and (D) 4-HO-TMT had no detectable levels when quantified via HPLC and thus suggested not to be capable of passively permeating a bilipid layer membrane.

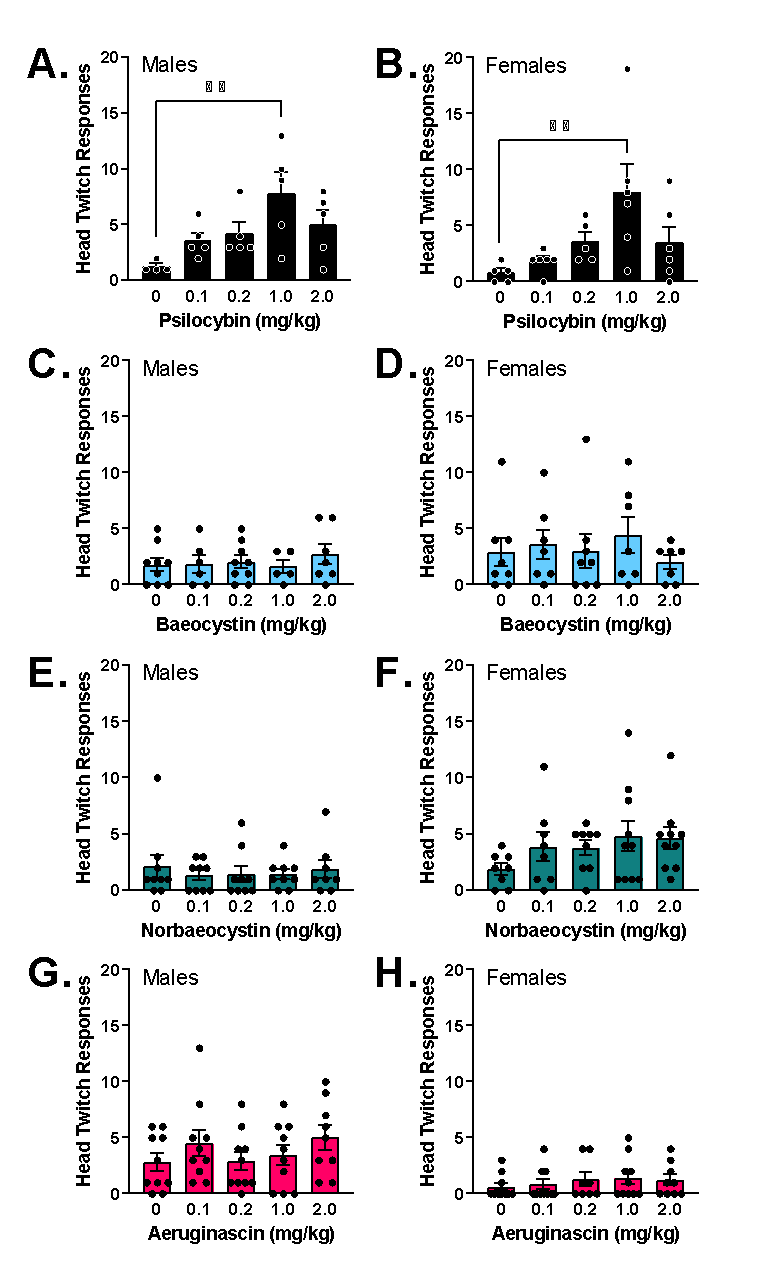

Supplemental Figure S6: Head twitch responses in animals receiving varying doses of tryptamines, separated by sex. Psilocybin in both females and males was demonstrated to significantly increase HTRs at a dose of 1mg/kg when compared with animals receiving vehicle. Combined sex data is presented as Figure 4 in the main text.

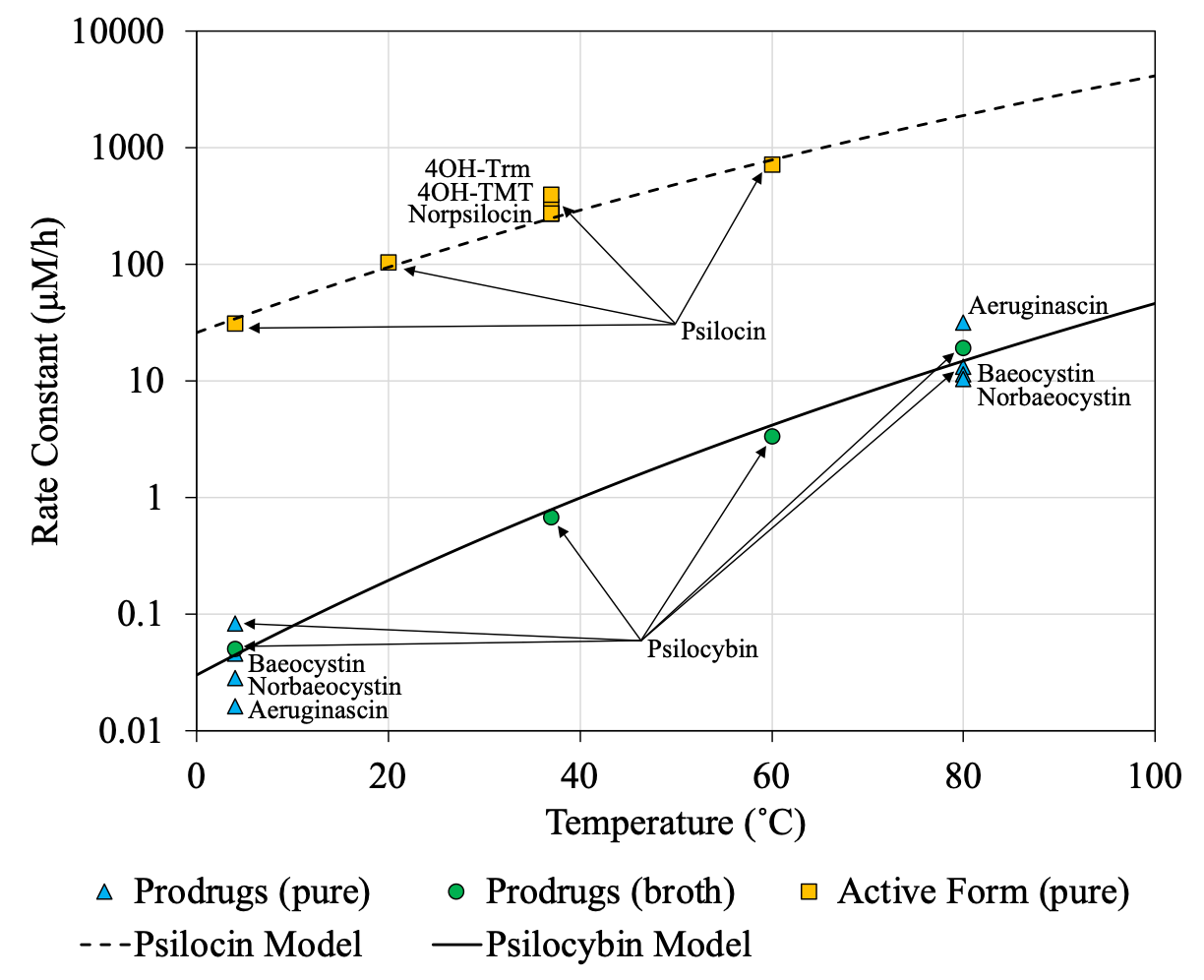

Supplemental Figure S7: Rate constants from zero order thermal degradation studies as a function of temperature for tryptamine prodrugs and dephosphorylated, active forms. Samples denoted as ‘pure’ were tested as purified drug in water, while those denoted as ‘broth’ were unpurified in spent, filtered *E. coli* cell broth. The solid and dashed lines represent the Arrhenius equation fitted to the psilocybin and psilocin data, respectively. All compounds had initial concentrations in the 3-5 mM range (approximately 1 mg/mL).

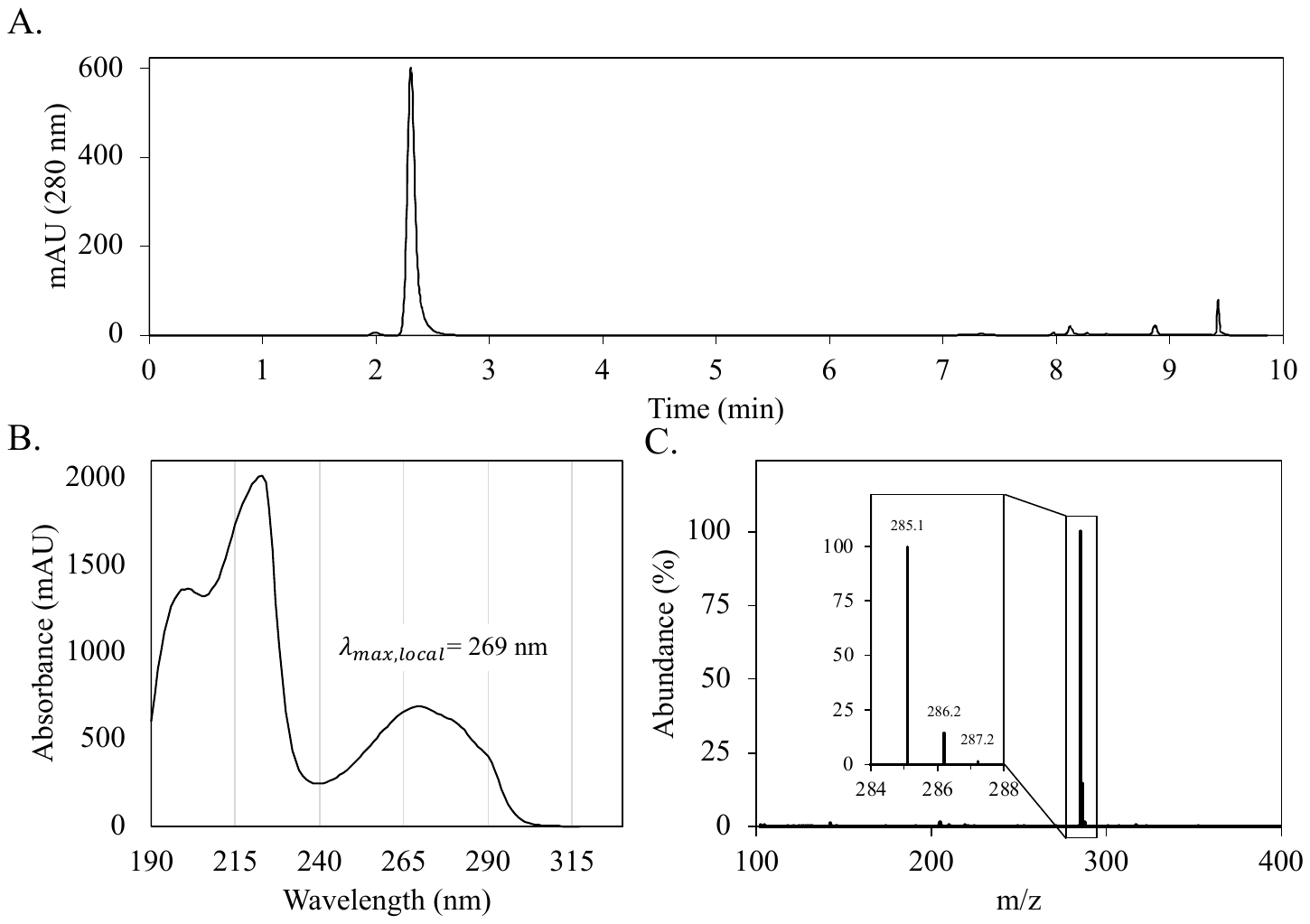

Supplementary Figure S8: Psilocybin characterization by (A) HPLC-UV at 280 nm, (B) UV absorption spectrum 190 nm – 330 nm, and (C) LCMS.

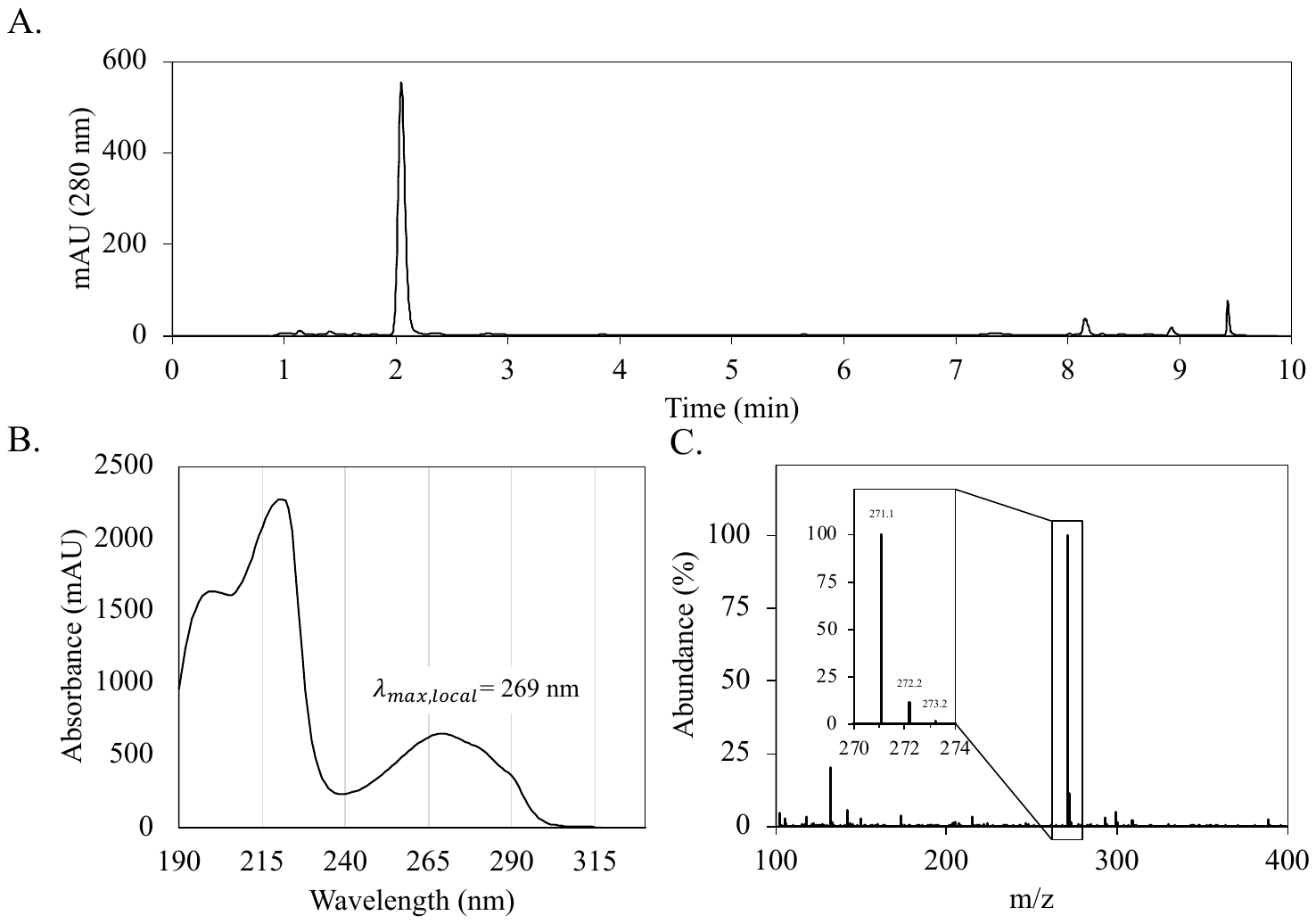

Supplementary Figure S9: Baeocystin characterization by (A) HPLC-UV at 280 nm, (B) UV absorption spectrum 190 nm – 330 nm, and (C) LCMS.

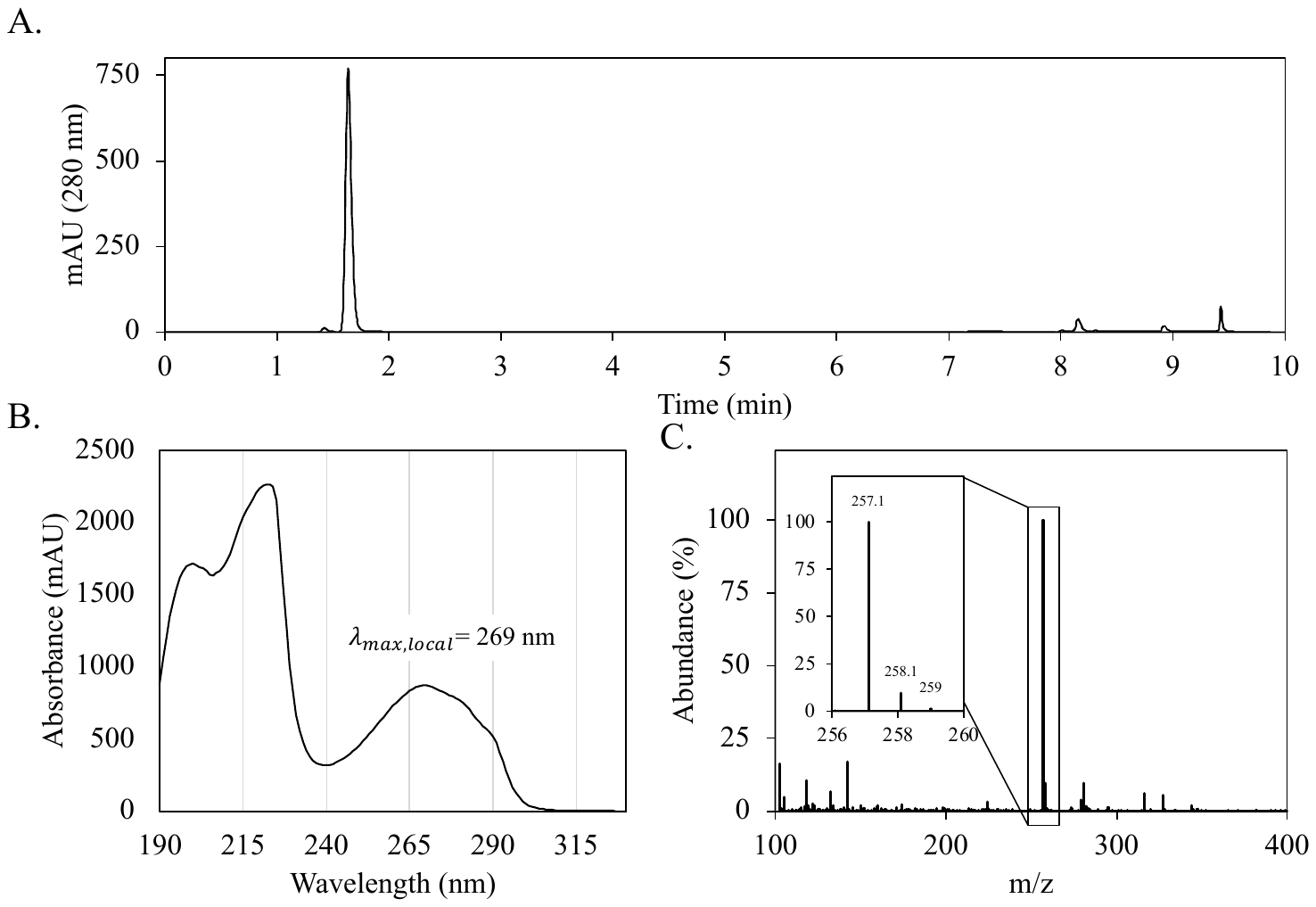

Supplementary Figure S10: Norbaeocystin characterization by (A) HPLC-UV at 280 nm, (B) UV absorption spectrum 190 nm – 330 nm, and (C) LCMS.

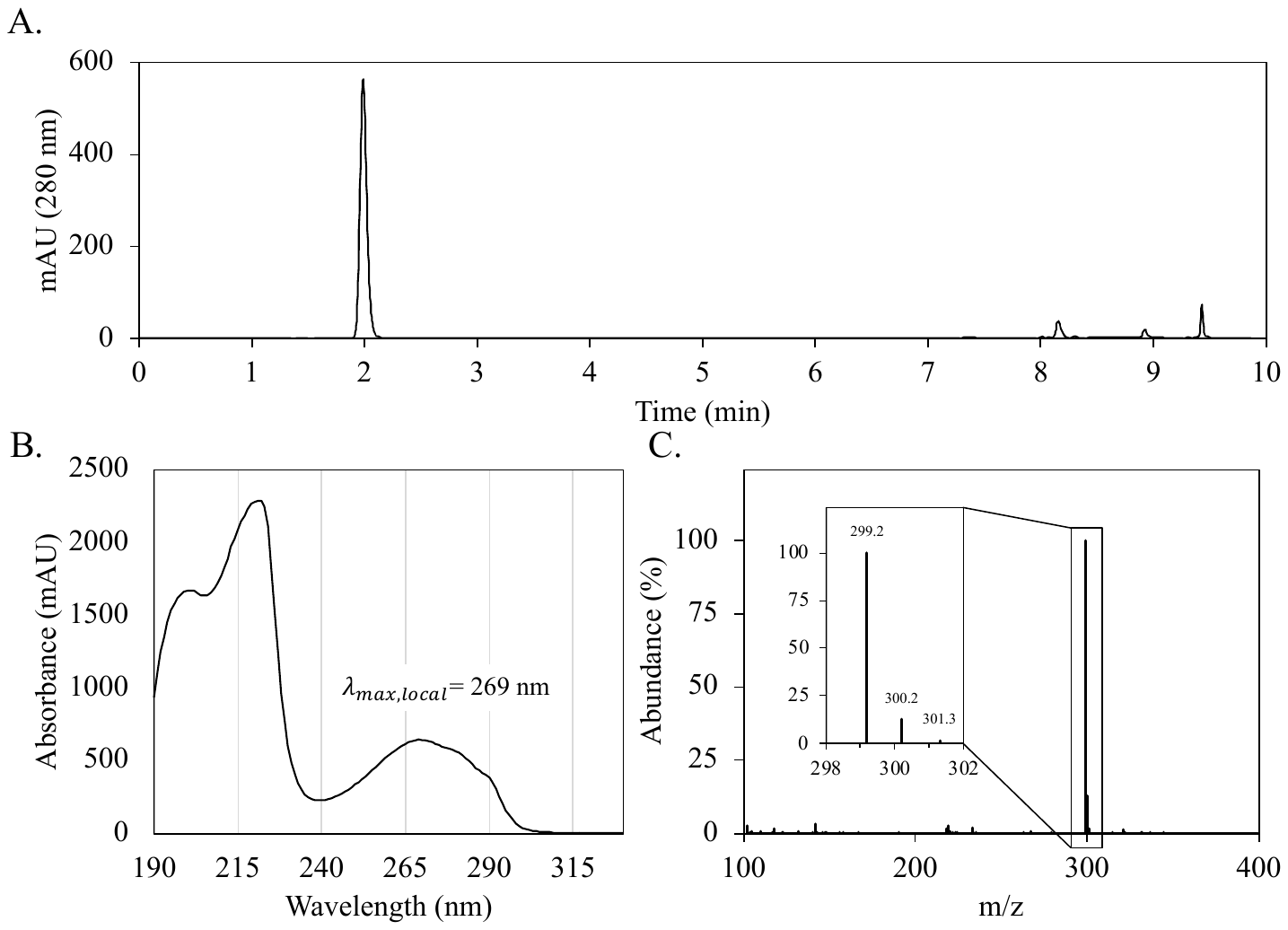

Supplementary Figure S11: Aeruginascin characterization by (A) HPLC-UV at 280 nm, (B) UV absorption spectrum 190 nm – 330 nm, and (C) LCMS.

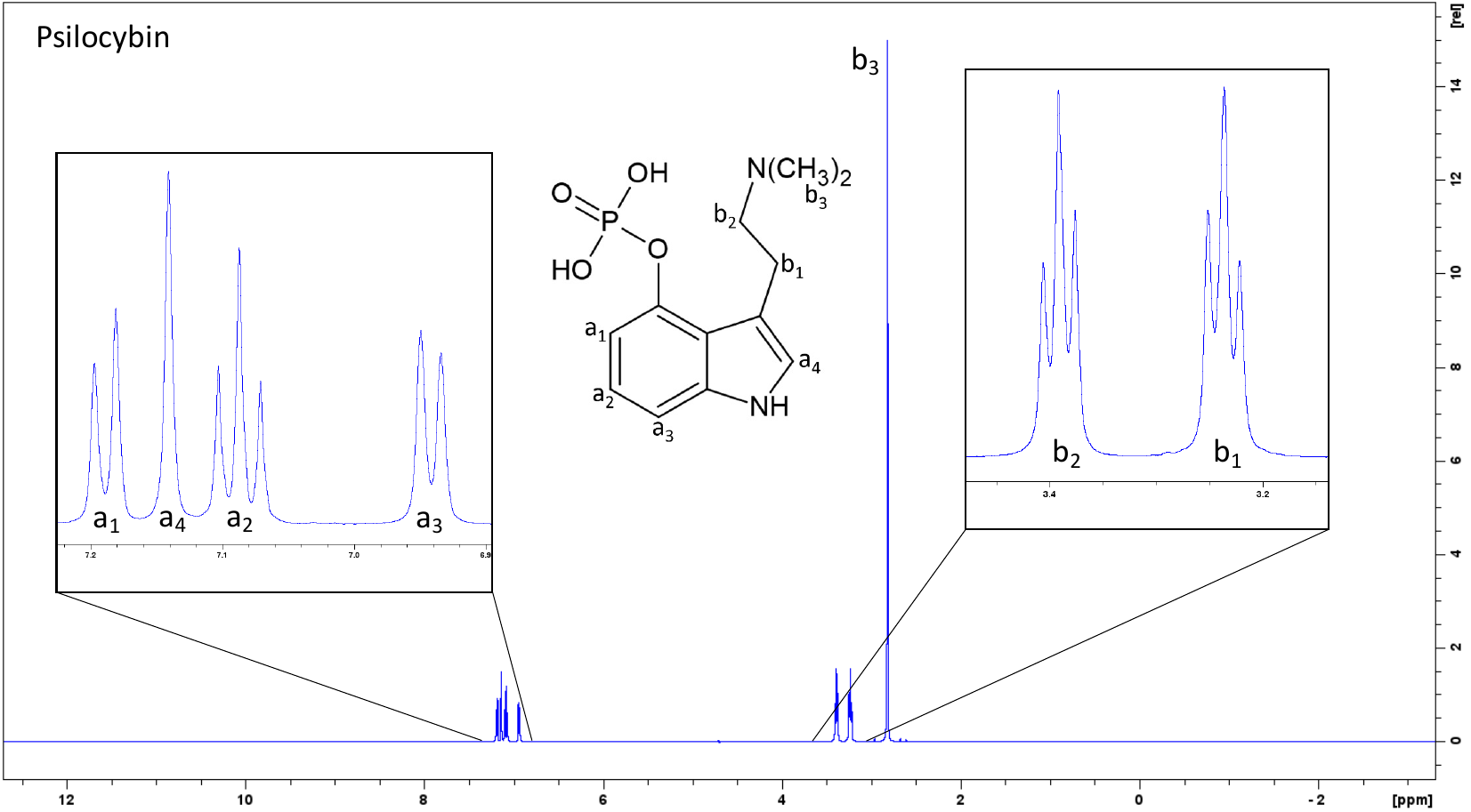

Supplementary Figure S12: Psilocybin ^1^H-NMR spectrum with peak assignments. ^1^H NMR (500 MHz, D_2_O) δ (ppm) 7.19 (1H, d, J = 8.1 Hz), 7.14 (1H, s), 7.09 (1H, t, J = 8.0 Hz), 6.94 (1H, d, J = 7.7 Hz), 3.39 (2H, t, J = 7.5 Hz), 3.24 (2H, t, J = 7.4 Hz), 2.82 (6H, s). Purity was determined to be 98.8%.

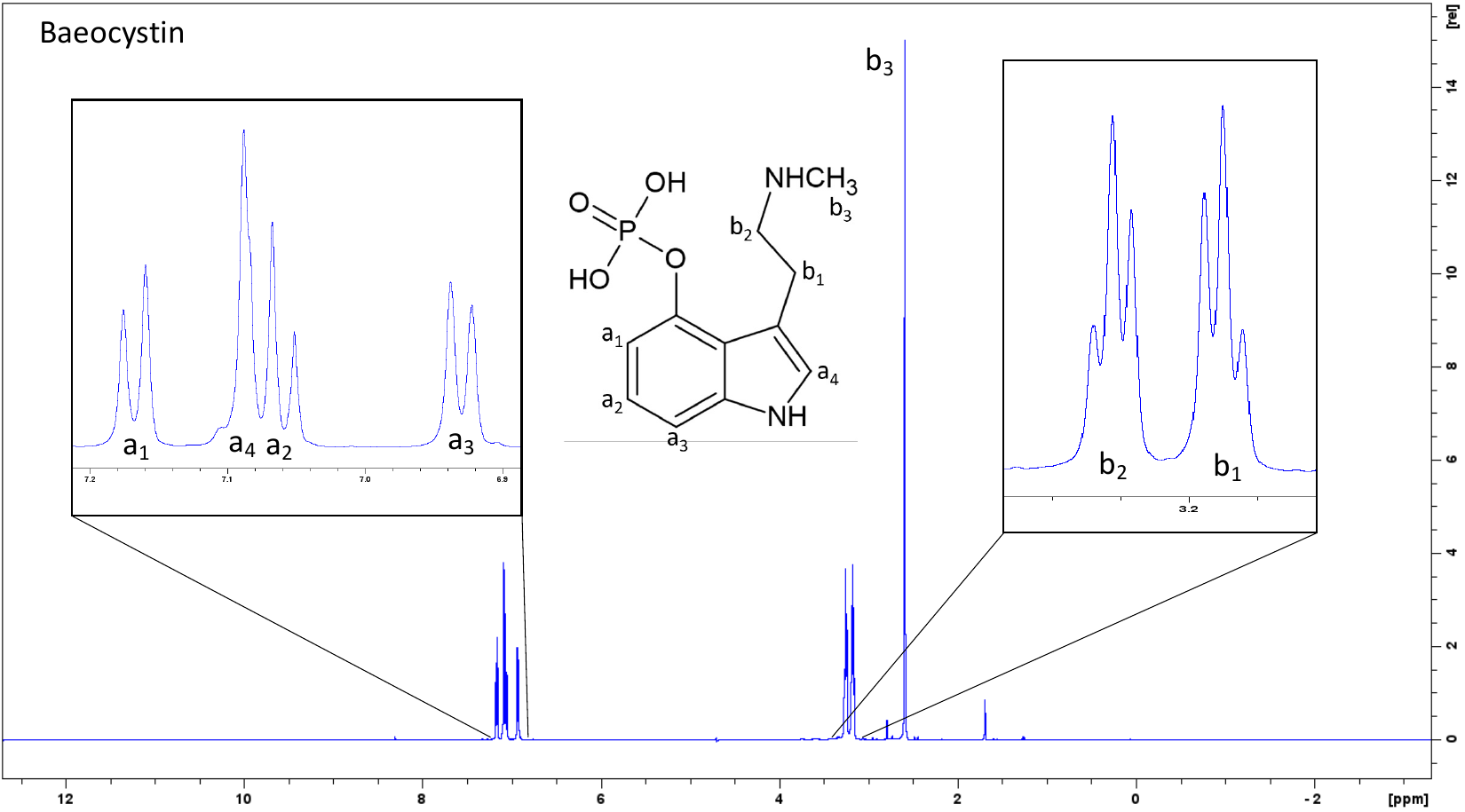

Supplementary Figure S13: Baeocystin ^1^H-NMR spectrum with peak assignments. ^1^H NMR (500 MHz, D_2_O) δ (ppm) 7.17 (1H, d, J = 8.2 Hz), 7.09 (1H, s), 7.06 (1H, m, J = 8.1 Hz), 6.93 (1H, d, J = 7.7 Hz), 3.26 (2H, t, J = 6.9 Hz), 3.17 (2H, t, J = 6.9 Hz), 2.60 (3H, s). Purity was determined to be 94.6%.

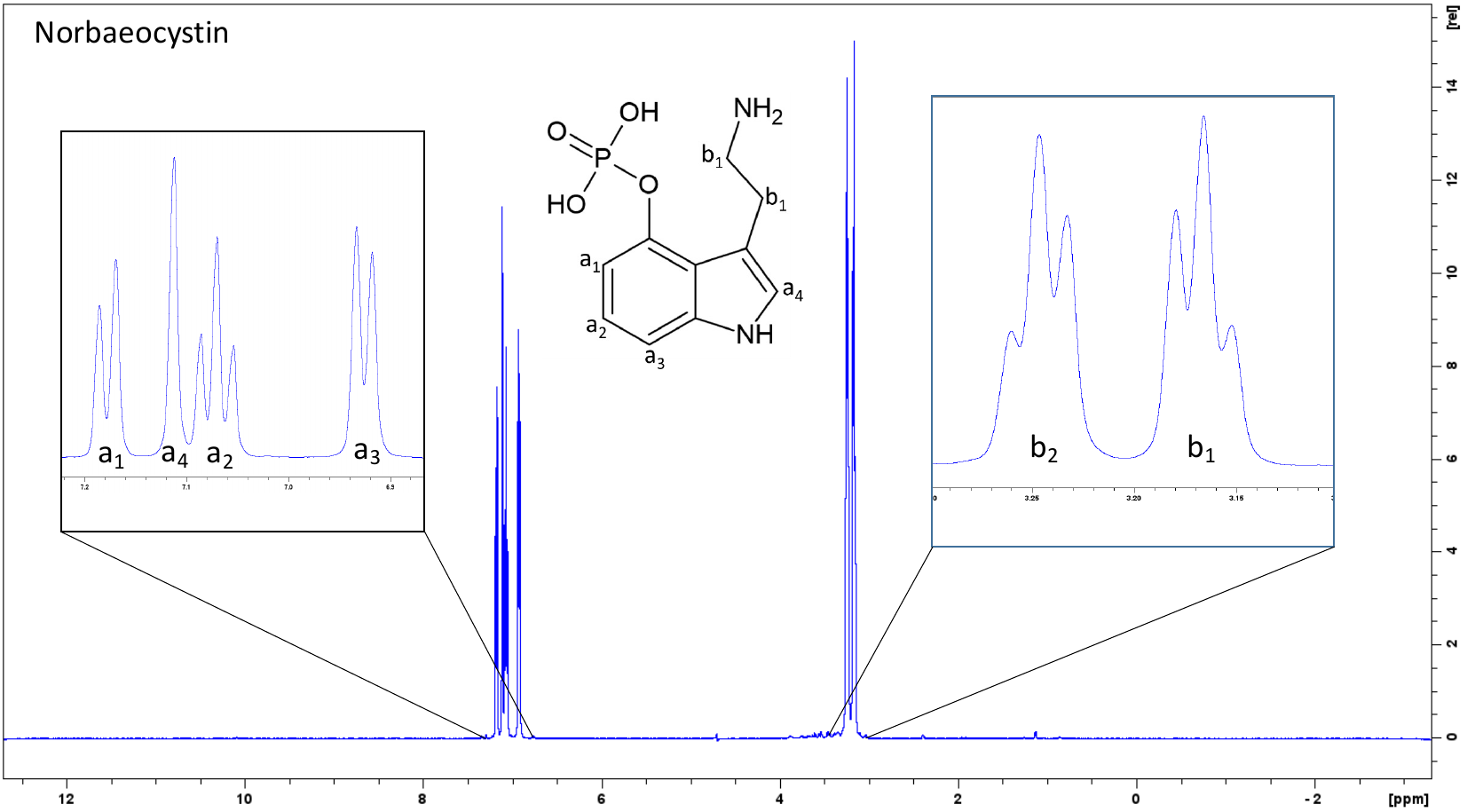

Supplementary Figure S14: Norbaeocystin ^1^H-NMR spectrum with peak assignments. ^1^H NMR (500 MHz, D_2_O) δ (ppm) 7.18 (1H, d, J = 8.1 Hz), 7.12 (1H, s), 7.07 (1H, t, J = 8.0 Hz), 6.92 (1H, d, J = 7.7 Hz), 3.25 (2H, t, J = 6.8 Hz), 3.16 (2H, t, J = 6.8 Hz). Purity was determined to be 97.6%.

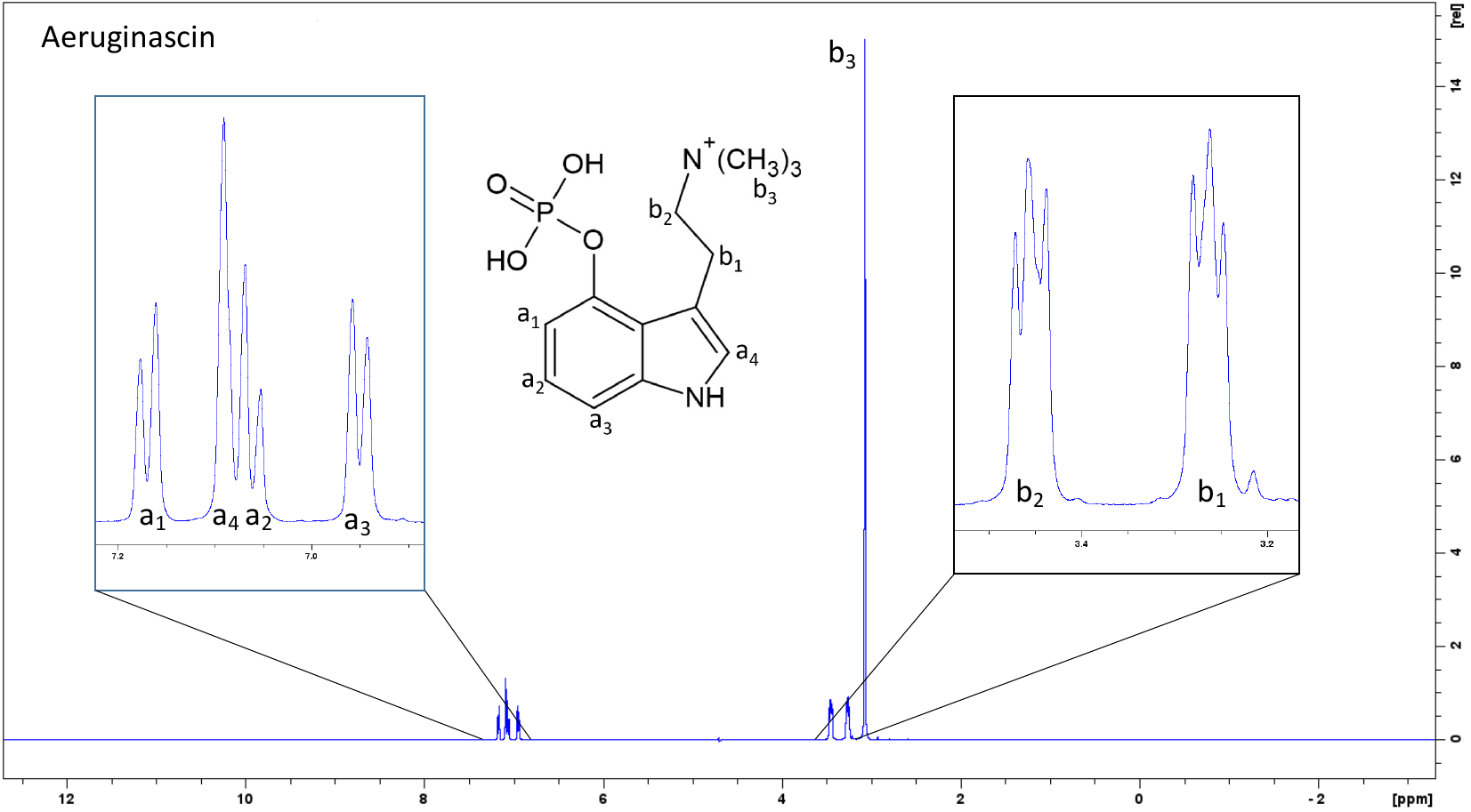

Supplementary Figure S15: Aeruginascin ^1^H-NMR spectrum with peak assignments. ^1^H NMR (500 MHz, D_2_O) δ (ppm) 7.17 (1H, d, J = 8.1 Hz), 7.09 (1H, s), 7.07 (1H, m, J = 8.1 Hz), 6.95 (1H, d, J = 7.6 Hz), 3.45 (2H, t, J = 8.3 Hz), 3.27 (2H, t, J = 8.2 Hz), 3.08 (9H, s). Purity was determined to be 98.6%.
